# Asymmetric inter-kingdom gene transfer and a horizontal transposon invasion in a powdery mildew fungus

**DOI:** 10.64898/2026.10.11.757921

**Authors:** Lee Mariault, Eric Lasserre, Jiangzhao Qian, Ji-Hun Hwang, Kyung Do Kim, Anaïs Gibert, Stefan Kusch, Ralph Panstruga, Moaine El Baidouri

## Abstract

Horizontal DNA transfer can introduce new genetic material into eukaryotic genomes, but how often recent transfers occur between multicellular lineages and what happens to transferred DNA after integration remain poorly understood. Plants and fungi provide a powerful system to address these questions because of their long-standing and intimate ecological associations. Here, we performed an annotation-independent, bidirectional whole-genome screen across 418 plant and 1,081 fungal genomes and identified 22 high-confidence recent horizontal transfer events. Twenty-one involved fungal DNA acquired by plants and were concentrated in bryophytes and Pooideae grasses, revealing pronounced directional and phylogenetic biases. The evolutionary fate of transferred DNA differed across recipient lineages. In Pooideae, fungal-derived genes underwent duplication, structural remodeling and expression divergence, and showed copy-number variation within species. In *Marchantia polymorpha*, transferred genes remained mostly single-copy but displayed presence–absence variation and ongoing loss among subspecies and natural accessions. In the opposite direction, we identified a plant-derived DNA transposon integrated into the genome of the tomato powdery mildew fungus *Pseudoidium neolycopersici*. Following transfer, this element remained active and generated a derived family of miniature inverted-repeat transposable elements that expanded to 172 copies. Recent detectable DNA transfer between plants and fungi is therefore uncommon and strongly asymmetric, but transferred sequences can remain highly dynamic after integration. Their subsequent loss, duplication, structural and regulatory diversification, or transposon-driven amplification links rare cross-kingdom acquisition events to ongoing genome variation and diversification.

## Background

Fungi and plants have interacted intimately for more than 450 million years, since the earliest colonization of terrestrial environments by embryophytes. Fossil evidence [1], together with phylogenomic and comparative genomic analyses [2, 3], indicates that symbiotic associations with fungi were already established in the earliest land plants [4] and likely played a key role in facilitating adaptation to terrestrial constraints such as nutrient limitation, desiccation, and fluctuating environmental conditions [5]. These interactions encompass a wide ecological spectrum, from mutualistic to pathogenic relationships, and involve both root and leaf tissues, either as endophytic or epiphytic associations.

Beyond their ecological and physiological importance, the persistence of close physical associations between plants and fungi over deep evolutionary timescales has also created conditions favorable for cross-kingdom horizontal transfer (HT), the non-sexual transfer of genetic material between unrelated organisms. Once considered rare in multicellular eukaryotes, HT is now recognized as a recurrent process that has contributed to the evolutionary history of land plants, particularly through transfers from microbial and fungal donors [6]. Large-scale phylogenomic analyses have identified two major ancient episodes of gene acquisition during early streptophyte evolution and the origin of land plants, during which hundreds of genes of bacterial and fungal origin were incorporated into plant genomes and retained in descendant lineages [7]. Many of these genes are involved in metabolism, stress responses, and interactions with the environment, suggesting that HT may have facilitated key evolutionary innovations during terrestrialization.

Recent studies further indicate that early-diverging land plants, particularly bryophytes, are especially enriched in horizontally acquired genes. A super-pangenome analysis across green plants revealed that bryophytes harbor a disproportionately large gene family space compared to vascular plants, partly driven by extensive acquisition of mainly microbial, but also, fungal genes through HT [8]. Pan-genome analysis of wild accessions of *Marchantia polymorpha* reveals that fungal fruit body lectin genes, acquired via horizontal gene transfer over 450 million years ago in the common ancestor of land plants, exhibit drought-induced upregulation, suggesting a functional role in abiotic stress tolerance [9].

Fungus-to-plant HT has also been documented in vascular plants, most prominently within grasses. Multiple independent studies have reported the acquisition of fungal genes from endophytic fungi of the genus *Epichloë* into cool-season grasses, including genes involved in cell wall modification, detoxification, and defense [10–12]. A striking example is the horizontally transferred glutathione S-transferase gene *Fhb7*, which confers resistance to *Fusarium* head blight in wheat relatives by detoxifying fungal mycotoxins [13]. These cases demonstrate that recent fungal-derived genes can be stably inherited, transcriptionally active, and functionally integrated into plant genomes.

Despite these advances, our understanding of the frequency, distribution, and molecular nature of recent inter-kingdom HT in plants remains incomplete. Many large-scale surveys relied heavily on annotation-based pipelines and protein-level similarity searches, which may underestimate the true extent of HT, particularly for recently transferred genes that remain unannotated, and/or for transfers involving non-coding regions [6]. Moreover, most studies are gene-centric, leaving open the question of whether larger genomic segments, including regulatory regions and introns, or transposable elements (TEs) are also transferred and how such sequences evolve following integration.

Here, we address these limitations by performing a whole-genome, annotation-independent screen for HTs between plants and fungi. By systematically comparing 418 plant genomes spanning the green lineage with 1,081 fungal genomes, and by combining stringent nucleotide-level similarity filters with phylogenetic reconstruction and manual inspection of genomic context, we identify 22 high-confidence recent inter-kingdom HT events. These include 21 fungus-to-plant transfers and a single, unambiguous case of plant-to-fungus transfer. The detected recent fungus-to-plant HT events are strikingly uneven in their phylogenetic distribution restricted mainly to bryophytes, and the Pooideae subfamily within angiosperms. In the latter, all detected transfers originate from *Epichloë* endophytes, reinforcing the view that tight and persistent endosymbioses promote recurrent gene exchange. In non-vascular plants, recent HT events are concentrated in bryophytes (particularly *M. polymorpha*) and predominantly originate from parasitic fungi of the order Pleosporales.

Our analysis further reveals marked contrasts in the post-transfer evolutionary trajectories of HT-derived genes across plant lineages. In Pooideae, several recently transferred genes have undergone tandem or dispersed duplication, resulting in divergent expression profiles, and in some cases retain signatures of DNA-mediated transfer, such as introns and extended flanking regions, followed by rapid structural remodeling. In non-vascular plants, by contrast, HT-derived genes are predominantly retained as single-copy loci, show comparatively stable expression across tissues and conditions, and exhibit limited retention of non-coding fungal sequence. Strikingly, we detect both signatures of recent HT gene acquisition and ongoing copy number variation and loss of HT-derived sequences in both Pooideae and bryophytes.

Finally, our genome-wide approach uncovers a rare and unique example of cross-kingdom transfer in the opposite direction: a pogo-like DNA transposon of plant origin, which we named *Gnawa* element, that has integrated into the genome of the powdery mildew fungus *Pseudoidium neolycopersici*, a pathogen of tomato, where it has been reactivated and massively amplified. Together, these findings provide a refined view of inter-kingdom DNA exchange, revealing its uneven distribution across plant lineages and highlighting how ecological context and contrasting post-transfer fates shape the evolutionary dynamics of transferred sequences in both plant and fungi genomes.

## Results

### Genome-wide screen for plant-fungus transfer

To characterize HT at the genome-wide scale without relying on prior genome annotation, we analyzed 418 plant genome assemblies obtained from the National Center for Biotechnology Information (NCBI), encompassing angiosperms (338), gymnosperms (6), lycophytes (3), bryophytes (5), monilophytes (2), charophytes (4), chlorophytes (52), and rhodophytes (8) (Additional file 1: Table S1), together with 1,081 fungal genomes from MycoCosm (Joint Genome Institute, JGI) and NCBI, primarily representing Ascomycota (66% of the dataset) and Basidiomycota (26%) (Additional file 1: Table S2). Pairwise genome-wide comparisons between fungal and plant genomes generated approximately 138 million conserved sequence matches from ∼400,000 whole-genome comparisons. To identify highly conserved sequences potentially resulting from HT while minimizing false positives, we retained only high-scoring pairs (HSPs) displaying ≥70% nucleotide identity over at least 500 bp and located on scaffolds larger than 20 kb in both fungal and plant genomes (see Methods). This initial filtering step reduced the dataset to 1.28 million candidate regions (Fig. 1).

**Fig 1.**
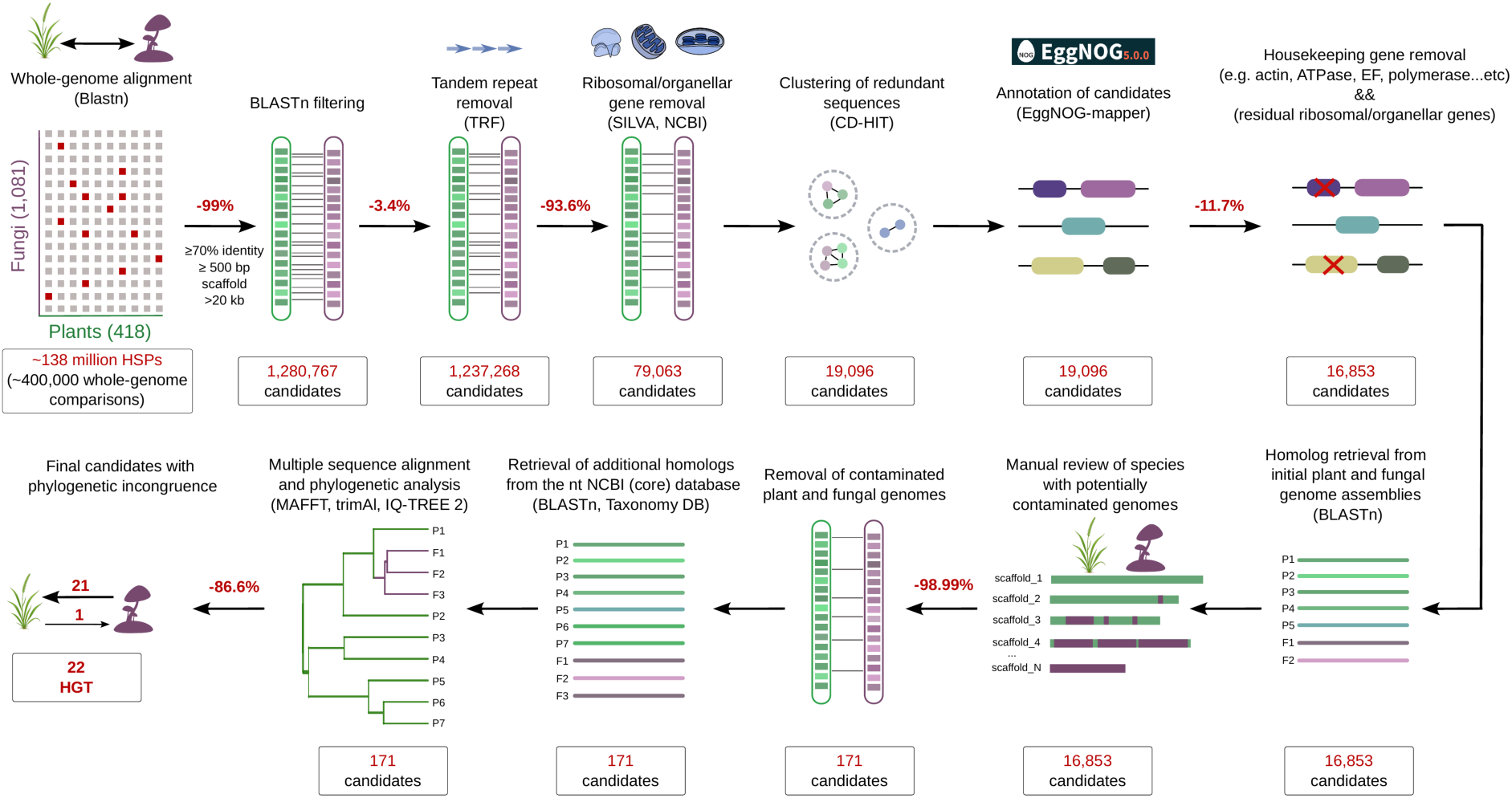
Overview of the bioinformatic pipeline for identifying horizontal transfer (HT) candidates between fungi and plants. Whole-genome BLASTn comparisons between 418 plant and 1,081 fungal genome assemblies were followed by successive filtering, manual curation of potential contaminants, homolog retrieval, and phylogenetic analyses to identify high-confidence HT events. Numbers indicate the number of candidate loci remaining after each filtering step, and percentages indicate the proportion of candidates removed. See the main text, Methods, and the GitHub repository [14] for further details.

The overwhelming majority of candidate regions corresponded to ribosomal, organellar, or housekeeping genes, whereas tandem repeats represented only a small fraction of the initial dataset. Successive filtering steps progressively reduced the number of candidate regions from approximately 1.3 million to ∼79,000 following the removal of tandem repeats and ribosomal/organellar sequences, and to ∼19,000 non-redundant loci after clustering. Annotation-based filtering further excluded housekeeping genes, leaving 16,853 candidate loci.

Most of these candidates were subsequently identified as likely assembly contaminants during manual inspection. Approximately 99% consisted of fungal-like sequences detected in only a few plant genome assemblies and typically confined to one or a few short scaffolds, a distribution inconsistent with stable genomic integration and indicative of contamination. Candidate loci were therefore examined in their genomic context and, whenever possible, compared with independently assembled genomes from the same or closely related species to assess their presence and exclude assembly artifacts. This curation step reduced the dataset to 171 candidate loci (Fig. 1).

For each remaining candidate, homologous sequences were first retrieved from the initial collection of plant and fungal genome assemblies and subsequently expanded by searching the core NCBI nt database to improve taxonomic sampling. Multiple sequence alignments and phylogenetic trees were then reconstructed, and candidates were retained only when phylogenetic analyses revealed strongly supported incongruence between plant and fungal lineages that was inconsistent with vertical inheritance.

Following manual evaluation of phylogenetic support, we identified 22 high-confidence horizontal transfer events: 21 were fungus-to-plant and one was plant-to-fungus (Additional file 2: Figs. S1–S13; sequences in Additional file 3). The fungus-to-plant HTs corresponded to genic regions representing 12 distinct gene families and one intergenic region (Additional file 1: Table S3; Additional file 2: Fig. S14). By contrast, the single plant-to-fungus HT event involved a TE. The fungus-to-plant HTs were unevenly distributed across the green lineage: 13 events occurred in non-vascular plants (10 in bryophytes and 3 in algae), whereas the remaining eight events involved the Pooideae subfamily within angiosperms. This pattern aligns with previous reports highlighting the uneven distribution of HT across plant lineages [15].

### Recent transfers are restricted to Pooideae and originate from *Epichloë*

In angiosperms analyzed, all detected fungus-to-plant HTs were confined to the core Pooideae, here understood as the clade comprising the tribes Poeae, Bromeae and Triticeae, and originated exclusively from endophytic fungi of the genus *Epichloë* (Fig. 2a; Additional file 1: Table S3). This restricted phylogenetic distribution is consistent with previous studies reporting multiple independent HT events in the Pooideae. Among the six HT events identified here, five are HTs corresponding to previously described events, whereas one involves a non-genic region and represents a novel HT event. Together, these results reinforce earlier observations of recurrent HT from *Epichloë* into Pooideae [10–13, 15].

**Fig 2.**
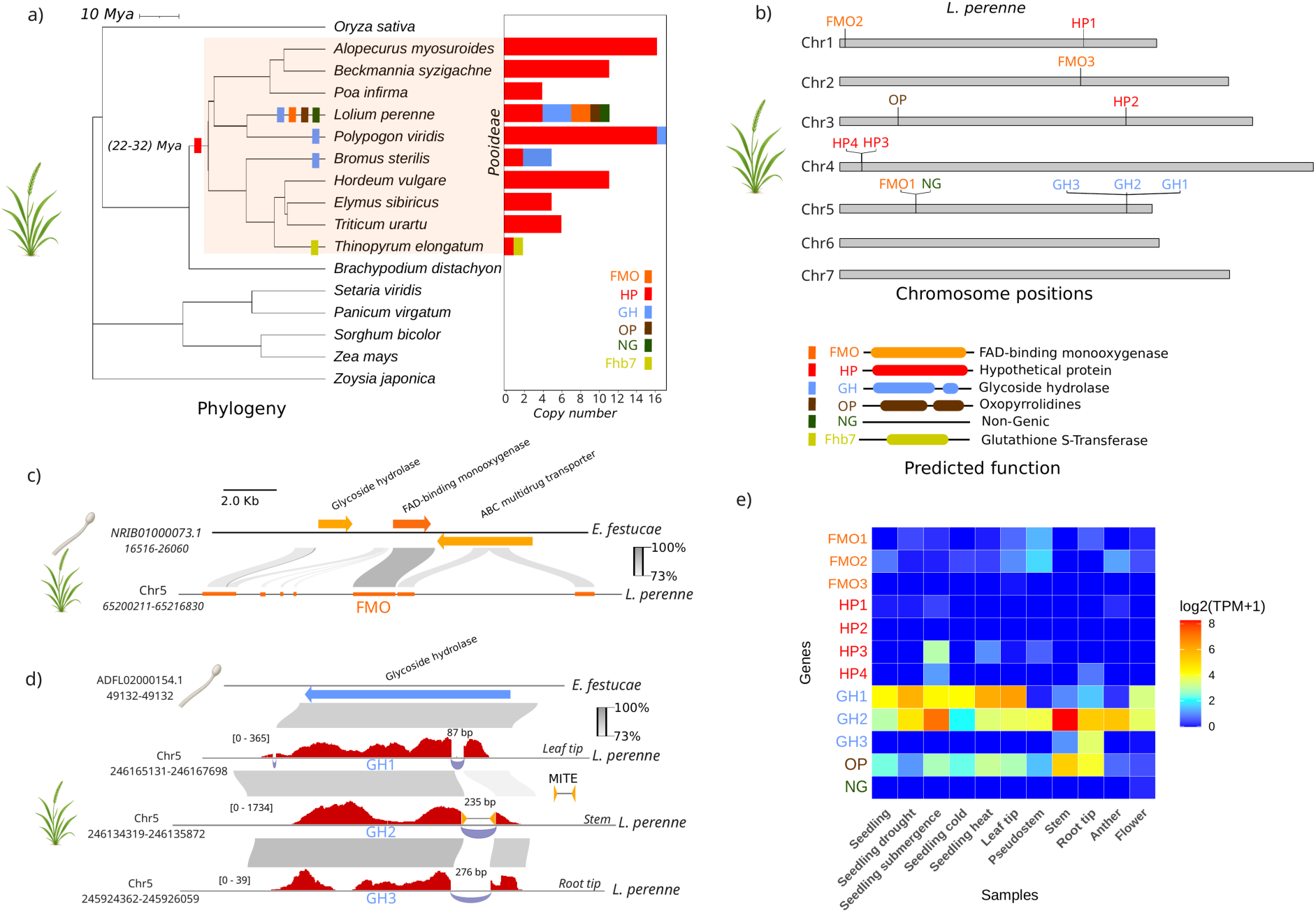
Horizontal gene transfer from *Epichloë* endophytes to Pooideae grasses. **a** Phylogenetic tree of Pooideae species and copy number variation of the six detected HT-derived genes (*FMO*, *HP*, *GH*, *OP*, *NG*, *Fhb7*). **b** Chromosomal distribution of the HT-derived paralogs within the *Lolium perenne* genome. **c** Synteny and large-block genomic transfer of the FAD-binding monooxygenase (FMO) locus between *Epichloë festucae* and *L. perenne*, illustrating the retention of extended flanking regions. Percentage represents sequence identity between *E. festucae* and *L. perenne*. **d** Post-transfer structural remodeling of the glycoside hydrolase (*GH*) paralogs in *L. perenne*, highlighting intron turnover and a MITE-derived insertion in *GH2*. **e** Expression profiles (log_2_(TPM + 1)) of the HT-derived genes across diverse tissues and stress conditions in *L. perenne* according to the color-coding of the heatmap shown on the right.

Within this clade, the six detected transfers resolve into two contrasting phylogenetic patterns: two exhibit broad distributions across multiple genera, whereas four are restricted to single lineages (Fig. 2a). The most widespread event involves a gene encoding a hypothetical protein (HP) of unknown function, which is present in all analyzed core Pooideae species, but is absent from *Brachypodium distachyon*, whose tribe is sister to the core Pooideae. This distribution is consistent with an ancient acquisition from an ancestral *Epichloë* lineage into the common ancestor of the core Pooideae, predating the diversification of this clade (Fig. 2a). Phylogenetic analysis of this gene further indicates that, in addition to this ancestral event, at least two independent and more recent transfers of the same gene have occurred between *Epichloë* and three Pooideae species: one into *Lolium perenne* and one shared by *Bromus sterilis* and *Elymus sibiricus* (Additional file 2: Fig. S2). Together, these patterns point to recurrent HT of the gene encoding HP within this host–symbiont system.

A second broadly distributed transfer involves a gene encoding a glycoside hydrolase (GH), which is detected in three genera (*Lolium*, *Polypogon*, and *Bromus*). This patchy distribution is consistent either with an early HT followed by multiple independent losses in other Pooideae lineages, or with recurrent, lineage-specific transfers from *Epichloë*. By contrast, four additional transfers are strictly lineage-specific. Notably, *L. perenne* harbors three unique HTs, including an oxopyrrolidines (*OP*) gene, a *FAD monooxygenase* (*FMO*), and a non-genic region (NG). Finally, we also recovered the previously described transfer in *Thinopyrum elongatum* encoding a *glutathione S-transferase*, *Fhb7* [13].

### Post-transfer duplication of fungus-derived genes in Pooideae

Copy-number profiling revealed that several HT-derived genes in Pooideae have undergone post-transfer duplication (Fig. 2a). Specifically, three of the five horizontally transferred genes show evidence of copy number expansion following their acquisition. The gene encoding the hypothetical protein (HP) has experienced multiple rounds of tandem duplication in nearly all Pooideae species, with copy numbers ranging from a single locus in *T. elongatum* to 16 copies in *Alopecurus myosuroides* (Fig. 2a,b; Additional file 2: Figs. S2 and S15).

A similar pattern was observed for the glycoside hydrolase (*GH*) gene, which underwent tandem duplication in both *L. perenne* and *B. sterilis* (Fig. 2b; Additional file 2: Figs. S3 and S15). In addition, the *FAD monooxygenase* (*FMO*) gene transferred to *L. perenne* has undergone dispersed duplication, giving rise to three distinct paralogs located on chromosomes 1, 2, and 5 (Fig. 2b; Additional file 2: Figs. S1 and S15). Together, these observations indicate that HT-derived genes in the Pooideae can undergo both tandem and dispersed duplication following their acquisition.

### Large genomic fragment transfer and subsequent remodelling in Pooideae

Genome-wide HT detection revealed that horizontally transferred sequences in the Pooideae frequently extend beyond annotated gene boundaries, indicating that some HT events involve the acquisition of larger genomic segments rather than isolated coding regions. A clear example is provided by the recent transfer of the *FMO* gene, which is embedded within a conserved genomic block that extends well beyond its coding sequence (Fig. 2c). Notably, we also detected an additional transfer of a non-genic region located approximately 38 kb downstream of the *FMO* locus (Fig. 2b). Together, these observations support a scenario in which the two *Epichloë*-derived sequences in *L. perenne* originated from a single horizontal transfer event encompassing a large genomic segment, followed by progressive erosion of non-functional sequences over evolutionary time. This scenario is further supported by sequence alignments between the *E. festucae* and *L. perenne* genomes, which show that nucleotide identity is highest across the *FMO* coding region, consistent with strong purifying selection on coding sequences, whereas the flanking regions exhibit markedly reduced conservation, suggesting relaxed selective constraints (Fig. 2c).

For the *glycoside hydrolase* (*GH*) gene, which comprises two exons, we found that the transfer includes the original fungal promoter region, and that one of the three *L. perenne* paralogs (*GH1*) retains the original 87-bp fungal intron, consistent with a DNA-mediated transfer rather than an RNA intermediate. By contrast, the duplicated paralogs show substantial post-transfer structural remodeling: in *GH2*, the fungal intron has been completely replaced by a 235-bp endogenous insertion derived from a Miniature Inverted-repeat Transposable Elements (MITE), whereas *GH3* has acquired a novel 276-bp intron of unknown origin, with no detectable similarity to either *Epichloë* or *L. perenne* sequences (Fig. 2d). These patterns illustrate rapid intron turnover and structural divergence following gene transfer.

The *OP* gene further illustrates differential retention of transferred sequence context. This gene shows detectable sequence conservation in the downstream region and within the intron, but no recognizable conservation in the promoter region (Additional file 2: Fig. S16), indicating partial retention of the original genomic environment. Finally, the transferred *HP* gene, which predates the diversification of the core Pooideae, shows no detectable sequence conservation beyond its coding boundaries (Additional file 2: Fig. S16), consistent with extensive erosion of non-coding regions following an ancient transfer.

### Expression divergence between transferred genes paralogs in Pooideae

To assess the expression pattern of horizontally acquired genes, we analyzed *L. perenne* transcriptomes spanning 11 tissues and stress conditions (Additional file 1: Table S4). Among the four transferred gene families examined, *GH* and *OP* showed moderate-to-high expression, whereas *FMO* and *HP* were largely transcriptionally silent (Fig. 2e). The *GH* locus displayed the strongest expression, reaching 306 transcripts per million (TPM) in stem tissue. Its three tandem paralogs exhibited marked regulatory divergence: *GH1* and *GH2* were broadly expressed, whereas *GH3* expression was restricted to root tips. The *OP* locus showed intermediate expression, peaking at 37 TPM in stem tissue. By contrast, *FMO* expression was negligible and reached only 2.2 TPM in pseudostem tissue. Despite its ancient acquisition and broad conservation across Pooideae, the *HP* family showed minimal expression across nearly all tissues and conditions. Only *HP3* was weakly expressed under submergence stress, reaching 6.7 TPM.

We next examined whether these expression patterns were conserved across Pooideae species. Analysis of leaf transcriptomes from eight species confirmed pervasive low expression of *HP* across most species and paralogs, with limited low-level expression detected only in *Beckmannia syzigachne* and *Elymus sibiricus* (Additional file 2: Fig. S17). In *Bromus sterilis*, two *GH* paralogs were expressed in leaf tissue, whereas a third remained undetectable. Finally, as expected, the transferred non-genic regions located on chromosome 5 of *L. perenne* showed no detectable transcriptional signal.

### Copy number variation of horizontally acquired loci in *Lolium perenne*

To explore intraspecific variation in horizontally acquired loci in Pooideae, we focused on *L. perenne*, which is the only species with two high-quality reference genomes available. We therefore investigated whether these HT-derived loci show copy number variation between the ‘Kyuss’ and ‘P226/135/16’ reference genomes [16, 17]. This analysis revealed two notable differences in HT copy number between the two genotypes (Additional file 2: Fig. S18). First, a paralog of the *FMO* gene located on chromosome 1 of ‘Kyuss’ is absent from the ‘P226/135/16’ reference. Conversely, the *HP* locus on chromosome 4, which is present as two tandem duplicates in ‘Kyuss’, includes an additional tandem copy in ‘P226/135/16’. In the absence of an appropriate outgroup to infer directionality, both gene loss through deletion and novel duplication events remain plausible explanations for these copy number differences.

### Bryophytes are enriched in horizontal gene transfers from Pleosporales

Genome-wide HT screening in non-vascular plants revealed a strong phylogenetic bias in both recipients and donors: recent fungus-to-plant transfers are overwhelmingly concentrated in bryophytes and predominantly originate from fungi of the order Pleosporales (Fig. 3; Additional file 2: Figs. S6–S12; Additional file 1: Table S3). In total, we identified 13 recent HT events involving seven distinct genes: ten events in bryophytes and three independent glutathione S-transferase (GST) transfers in green algae (Fig. 3). Within bryophytes, six events were detected in *Marchantia polymorpha*, two in *Ceratodon purpureus*, and one each in *Lunularia cruciata*, *Physcomitrium patens*, and *Syntrichia caninervis*. One HT event was shared by *C. purpureus* and *S. caninervis* and was therefore counted as a single ancestral transfer event (Additional file 2: Fig. S11). Notably, only two of the 13 HT events had been reported previously [8, 15], indicating that the majority of detected transfers represent previously unreported HTs (Additional file 1: Table S3).

**Fig 3.**
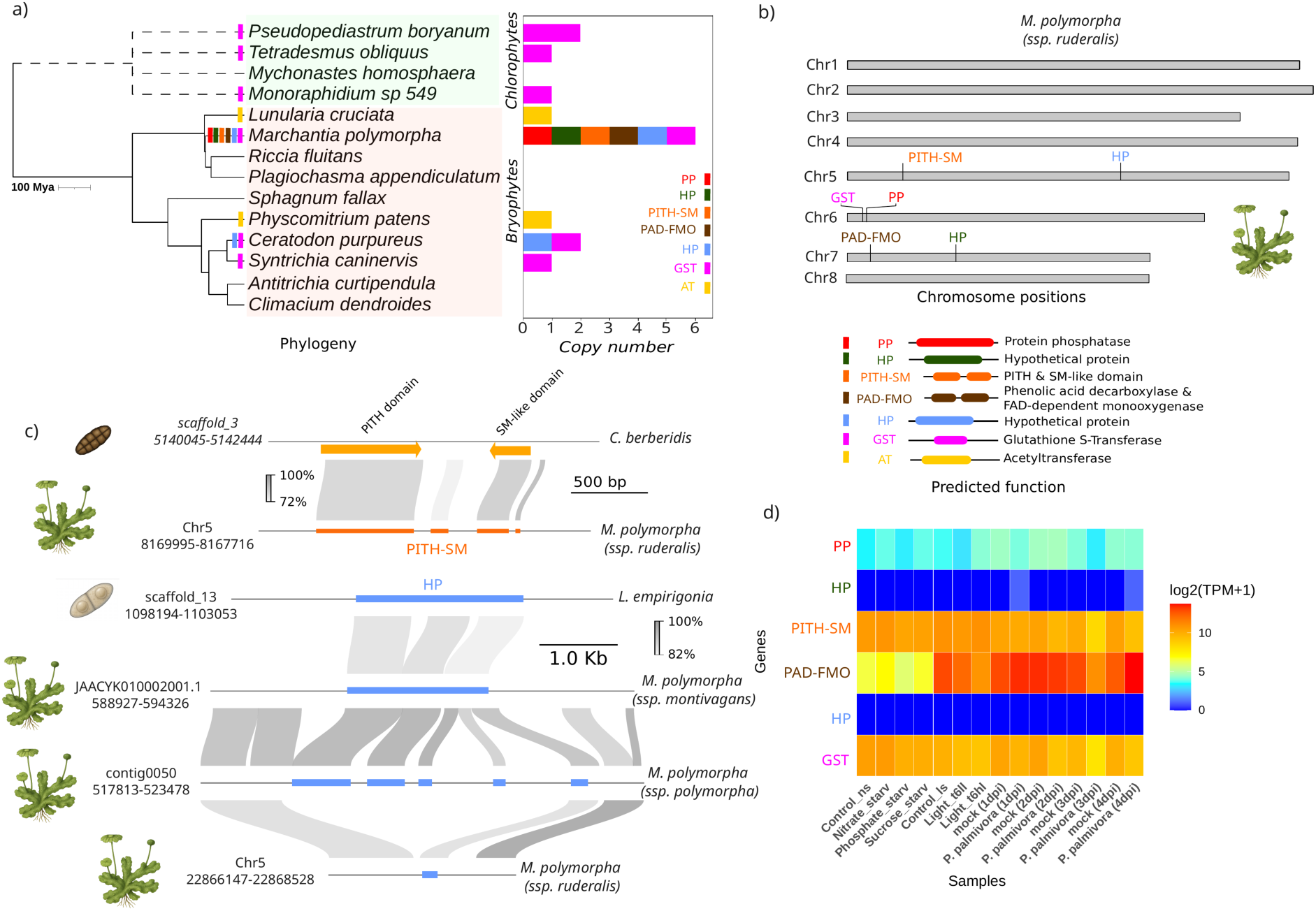
Horizontal gene transfer in non-vascular plants (bryophytes and green algae). **a** Phylogenetic tree of green algae and bryophytes and copy number variation of the horizontally acquired genes. **b** Chromosomal distribution of the HT-derived genes in the *Marchantia polymorpha* (subsp. *ruderalis*) genome. **c** Genomic alignments of representative HT loci between *M. polymorpha* and homologous regions of closely related fungal genomes (*Cucurbitaria berberidis* for *PITH-SM* and *Lizonia empirigonia* for *HP*; schematically depicted on the left), together with comparisons across *M. polymorpha* subspecies, wild accessions, and related species, illustrating locus conservation and presence–absence polymorphism. Percentages indicate sequence identity between homologous regions. **d** Heatmap showing expression profiles (log_2_(TPM + 1)) of the transcriptionally active HT-derived genes across diverse experimental conditions.

The transferred genes encode a diverse functional repertoire, including two hypothetical proteins (HPs), a protein phosphatase (PP), a PITH & SM-like domain-containing protein (PITH-SM), a protein harboring phenolic acid decarboxylase and FAD-dependent monooxygenase domains (PAD-FMO), a glutathione S-transferase (GST), and an acetyltransferase (AT) (Fig. 3a). Despite this functional diversity, three genes (*GST*, *AT*, and *HP*) have undergone multiple independent HT events (Additional file 2: Figs. S10–S12), indicating recurrent transferability. In particular, the *GST* gene was transferred five times independently from fungi to both bryophytes and chlorophytes (Fig. 3a). The *HP* gene underwent independent transfers to *M. polymorpha* and *C. purpureus*, whereas the acetyltransferase was independently acquired by *L. cruciata* and *P. patens*.

For seven of the 13 HT events, phylogenetic analyses enabled inference of putative fungal donors (Additional file 2: Figs. S6–S12; Additional file 1: Table S3). Strikingly, six of these putative donors (*Fenestella fenestrata, Cucurbitaria berberidis, Corynespora cassiicola, Clathrospora elynae, Trematosphaeria pertusa,* and *Setomelanomma holmii*) belong to the order Pleosporales, a group encompassing saprotrophic, endophytic, and pathogenic fungi frequently associated with living or senescent plant tissues. The only exception involved the *HP* and *PAD-FMO* genes in *M. polymorpha*, which appears to derive respectively from *Coccomyces strobi* (Rhytismatales) and *Delphinella strobiligena* (Dothideales).

### Absence of post-transfer duplications in non-vascular plants

In contrast to the Pooideae (Fig. 2), HTs identified in non-vascular plants are predominantly maintained as single-copy loci, with no evidence for tandem or dispersed duplications (Fig. 3a). The only exception is the *GST* gene, which underwent a tandem duplication in the chlorophyte *Pseudopediastrum boryanum*, giving rise to two adjacent copies.

To evaluate whether these HT-derived genes are transcriptionally active in recipient lineages, we analysed publicly available transcriptomic datasets from bryophytes and chlorophytes harbouring at least one inferred HT event (Fig. 3a). These datasets encompass multiple tissues and stress conditions (Fig. 3d; Additional file 1: Table S4). In *M. polymorpha*, four of the six HT-derived genes are transcriptionally active with variant expression levels. While *PP* is expressed at low levels, with an average TPM of 1.22, *GST* and *PITH-SM* genes show moderate expression with average TPM values of 62.3 and 50.8, respectively. Notably, *PAD-FMO* exhibits the highest transcript abundance among HT derived genes in *M. polymorpha* with an average TPM of 217.8 (Fig. 3d). By contrast, the two hypothetical protein genes (*HPs*) show no detectable expression across the tissues and conditions analysed, mirroring the absence of expression observed for the *HP* gene horizontally transferred into *L. perenne*.

In the other bryophyte species and chlorophytes, with the exception of *P. boryanum*, HT-derived genes display moderate to high expression levels across diverse conditions (Additional file 2: Fig. S19). These include the *GST* gene in *S. caninervis*, *Tetradesmus obliquus* and *C. purpureus*, the *AT* gene in *P. patens* and *Pellia cruciata*, and the *HP* gene in *C. purpureus*. Notably, unlike HT-derived genes in the Pooideae, which frequently display condition-dependent expression variation associated with gene duplication, HT-derived genes in non-vascular plants exhibit comparatively stable expression profiles across tissues and stress treatments, consistent with their retention as single-copy loci.

### Limited retention of fungal non-coding sequence following transfer

In contrast to the extensive large-block transfers observed between *Epichloë* and *L. perenne*, we found no evidence for the horizontal acquisition of large genomic regions in non-vascular plants. Instead, recent HT events in bryophytes and chlorophytes appear to be largely restricted to coding sequences, with only limited retention of flanking non-coding DNA. Although the *GST* and *PP* genes in *M. polymorpha* are located near the beginning of chromosome 6 and are separated by only approximately 1 kb (Fig. 3b), their inferred origins from distinct fungal donors (Additional file 1: Table S3) argue against a single large-block transfer event. Instead, these loci most likely represent independent acquisition events that fortuitously inserted into nearby genomic positions, which may represent “hotspots” for the insertion of horizontally transferred DNA.

The main exception involves the *PAD-FMO* gene in *M. polymorpha*, for which RNA-seq data indicate that the promoter region may also be of fungal origin, extending up to 52 bp upstream of the transcription start site (Additional file 2: Fig. S20a). Finally, the *PITH-SM* gene in *M. polymorpha* is embedded within a larger endogenous gene (MpTK2_5g06260) (Additional file 2: Fig. S20b). This locus comprises two major exons, indicating that horizontally acquired sequences can be co-opted into pre-existing gene architectures and contribute to the formation of chimeric genes, thereby participating in genomic plasticity.

### Horizontal gene transfer variation across *Marchantia polymorpha* subspecies

*M. polymorpha* comprises three major subspecies (*M. polymorpha* ssp. *montivagans*, ssp. *polymorpha*, and ssp. *ruderalis*), with ssp. *montivagans* forming an outgroup relative to ssp. *polymorpha* and ssp. *ruderalis* [9]. Genome-wide HT detection using the reference genome of these subspecies revealed clear differences in HT content among them. Of the six HT events identified in *M. polymorpha*, three fungal-derived loci exhibit presence–absence polymorphisms and signatures of ongoing gene loss. As shown in Fig. 3c, the *HP* gene acquired from *Clathrospora elynae* is retained in ssp. *montivagans*, whereas the corresponding locus in ssp. *polymorpha* has undergone extensive structural variation, including multiple insertions that disrupt the coding sequence. In ssp. *ruderalis*, this *HP* locus is largely deleted, with only approximately 200 bp of the original fungal sequence remaining. The *PP* gene is detected in ssp. *polymorpha* and ssp. *ruderalis* but is absent from the outgroup ssp. *montivagans*, a pattern consistent either with a transfer event in the common ancestor of ssp. *polymorpha* and ssp. *ruderalis* or with secondary loss in ssp. *montivagans*. By contrast, the *PAD-FMO* gene is exclusively present in ssp. *ruderalis* and absent from both ssp. *montivagans* and ssp. *polymorpha*, consistent with a recent and lineage-specific HT event.

To exclude the possibility that these differences reflect assembly artifacts, we aligned whole-genome sequences from three natural wild accessions [9] per subspecies (NILSC12, RES33, and NILSC24 for ssp. *montivagans*; Caz-A, NILSC1, and WRG for ssp. *polymorpha*; and Ale-A, TAK1, and RES34 for ssp. *ruderalis*) to the reference genome of ssp. *ruderalis* genomes that have retained the six HT events (Additional file 1: Table S5). These comparisons fully confirm the presence–absence patterns inferred from the reference assemblies, supporting the conclusion that the detected HT variations within *M. polymorpha* reflect genuine biological polymorphisms in the content of horizontally transferred genes (Additional file 2: Fig. S21).

### A rare plant-to-fungus horizontal transfer in the tomato powdery mildew

Among the horizontal transfer events uncovered through our genome-wide comparisons, one is particularly notable: a plant-to-fungus transfer involving a ∼2.2 kb Class II DNA transposon in *P. neolycopersici*, the causal agent of tomato powdery mildew. We designate this new element *Gnawa*. No *Gnawa* homolog was detected in the tomato genome, in the 1,081 fungal genomes screened, or in raw sequencing reads from the related powdery mildews *Erysiphe alphitoides* (SRR10956459), *E. pisi* (SRR11059780) and *E. necator* (SRR18712276), supporting a recent lineage-specific acquisition. Sequence analysis indicates that *Gnawa* belongs to the *Tc1*/*mariner* superfamily and encodes the canonical domains characteristic of this group, including a centromere protein B–like domain, a *Tc5* transposase-binding domain, and a DDE superfamily endonuclease, consistent with an autonomous DNA transposon architecture (Fig. 4a, b). The element is flanked by 28-bp terminal inverted repeats (TIR). Sequence comparison and phylogenetic reconstruction based on the transposase and DDE domains of known *Tc1*/*mariner* elements [18] places *Gnawa* within the *Pogo* clade (Additional file 2: Fig. S22). Consistent with this affiliation, the catalytic core of the *Gnawa* transposase contains a DD31D motif, a signature shared with *mariner*-like and *Pogo*-like elements rather than *Tc1*-like elements (Fig. 4b). In total, we identified nine full-length copies of *Gnawa* in the *P. neolycopersici* genome, indicating post-transfer amplification (Fig. 4c).

**Fig 4.**
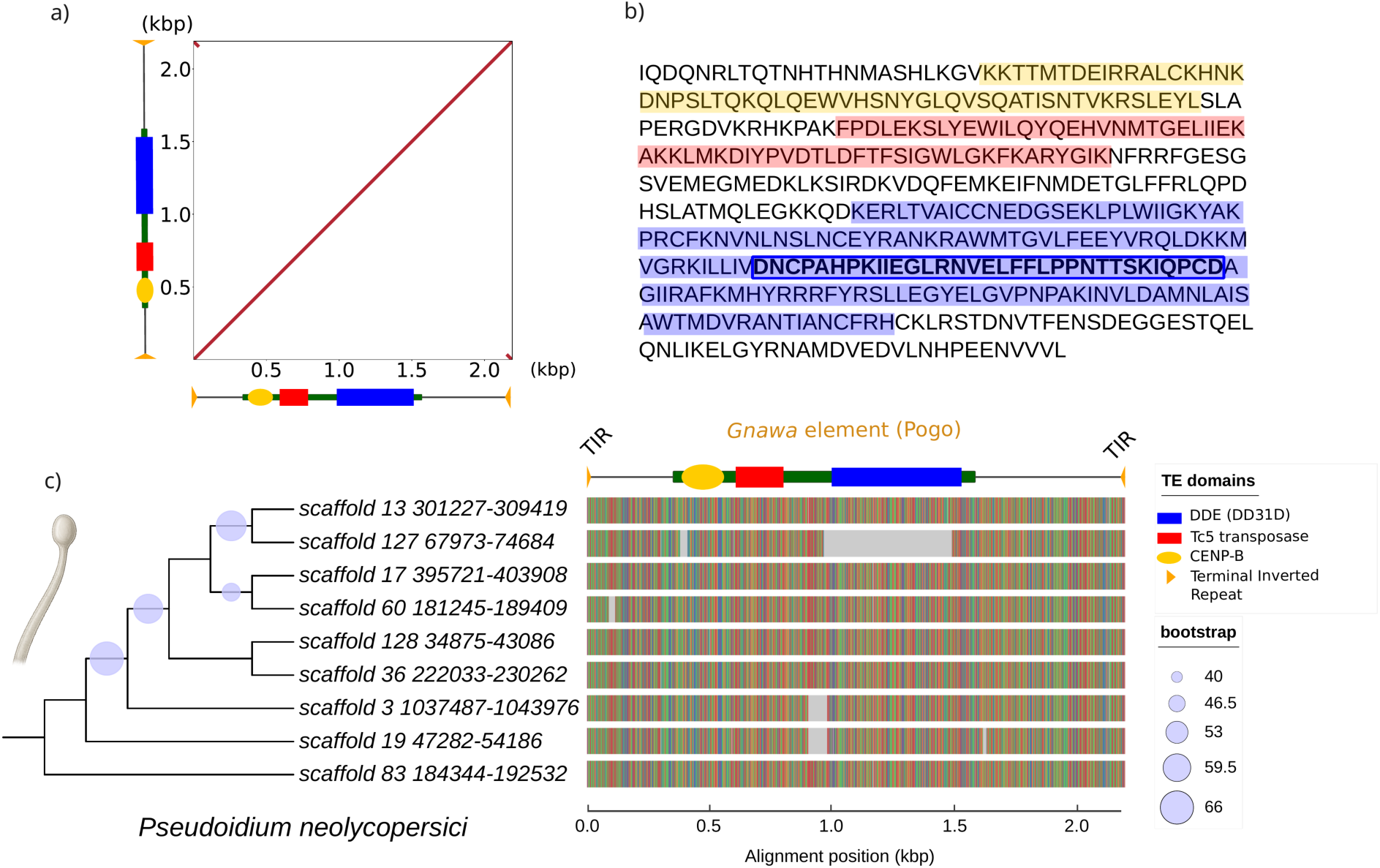
Structural organization and phylogeny of *Gnawa*, a plant-derived pogo transposon horizontally acquired by *Pseudoidium neolycopersici*. **a** Dot plot and structural organization of the 2.2-kb autonomous *Gnawa* element, showing its terminal inverted repeats (TIRs) and coding region. **b** Predicted amino acid sequence of the *Gnawa* transposase, highlighting the conserved CENP-B DNA-binding domain (yellow), *Tc5* transposase domain (red), and catalytic DDE (DD31D) motif (blue) characteristic of *Pogo* transposons. **c** Maximum-likelihood phylogeny and sequence alignment of the nine full-length autonomous *Gnawa* copies identified across the *P. neolycopersici* genome, showing the conservation of coding and non-coding regions, including the TIRs.

Homology searches recovered only plant-derived sequences but no homolog of this element has been found in the tomato genome. Consistent with a recent and lineage-specific acquisition, we detected no *Gnawa* element in the 1,081 screened fungal genomes. In addition, targeted searches of both reference genome assemblies and unassembled Illumina and PacBio sequencing reads from closely related powdery mildew species infecting oak (*Erysiphe alphitoides*), pea (*Erysiphe pisi*) and grapevine (*Erysiphe necator*) revealed no evidence of any *Gnawa* element.

To exclude the possibility that *Gnawa* represents plant DNA contamination or an assembly artefact, we systematically searched for the element in unassembled raw sequencing data from two independent *P. neolycopersici* strains sequenced using both PacBio and Illumina platforms (Additional file 2: Fig. S23). Whole-read mapping against the reference *Gnawa* sequences consistently detected the element in both datasets. Moreover, for each of the nine full-length copies, PacBio reads spanning the entire transposon and its flanking genomic regions were recovered, unambiguously demonstrating physical integration of *Gnawa* within the fungal genome (Additional file 2: Fig. S24). Together, these results rule out contamination and confirm that *Gnawa* represents a bona fide plant-derived transposon that has been horizontally acquired and stably integrated into the genome of *P. neolycopersici*.

### Recurrent transfer of *Gnawa* element across eudicots

Across eudicots, full-length (∼2.2 kb) *Gnawa* homologs exhibit a patchy distribution and frequent topological conflicts between element and host phylogenies (Fig. 5a,b), a pattern indicative of multiple transfer events. Notably, these homologs also display exceptionally high nucleotide identity and short branch lengths across distantly related plant lineages (Fig. 5d; Additional file 2: Figs. S13 and S25). For instance, *Gnawa* elements from *P. neolycopersici* share ∼99% nucleotide identity with homologs from Lamiales (*Veronica anagallis-aquatica*), Saxifragales (*Saxifraga aizoides*), Ericales (*Primula vulgaris* and *P. scotica*), Asterales (*Lophiolepis eriophora* and *Helianthus annuus*), and Malpighiales (*Viola palustris*, *V. epipsila*, and *V. suecica*) (Additional file 2: Fig. S25). In several of these species, within-genome divergence among *Gnawa* paralogs is minimal, and copy numbers can exceed 100 (e.g., in *V. suecica* and *V. epipsila*), consistent with recent transpositional activity and lineage-specific expansion following transfer (Additional file 2: Fig. S26). Phylogenetic reconstruction using full-length, non-redundant *Gnawa* copies places the *P. neolycopersici* sequences within the eudicot clade, forming a well-supported cluster (Fig. 5a,b). Among the analyzed plants, the closest homologs to the fungal copies are from *V. anagallis-aquatica* (Lamiales) and *Myriophyllum alterniflorum* (Saxifragales), with bootstrap support exceeding 70%.

**Fig 5.**
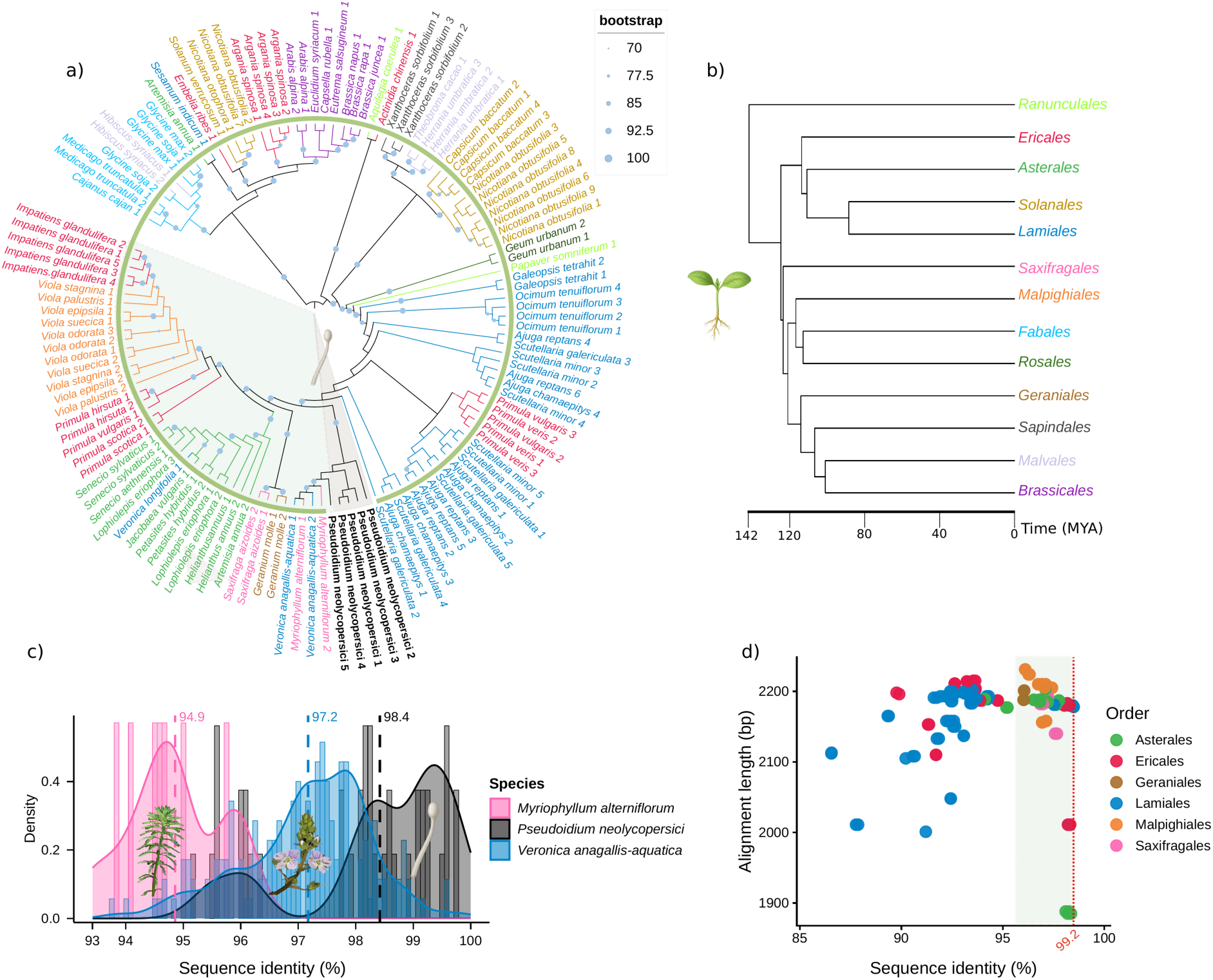
Widespread but patchy distribution of *Gnawa* homologs across eudicots. **a** Phylogenetic tree of full-length *Gnawa* copies across various dicot species, demonstrating incongruence with the host phylogeny. **b** Simplified timetree of the corresponding host plant orders. **c** Density distribution of intra-genomic paralog sequence identity (%) for *P. neolycopersici*, *Veronica anagallis-aquatica*, and *Myriophyllum alterniflorum*, used as a proxy to infer the relative timing of transpositional bursts. **d** Scatter plot of alignment length (bp) versus sequence identity (%) of *Gnawa* homologs, colored by host plant order.

To explore, in a tentative manner, the direction of *Gnawa* transfer between plants and *P. neolycopersici*, we analysed within-genome paralog identity as a proxy for relative age (Fig. 5c). This approach enables comparison of the relative timing of *Gnawa* transpositional activity among lineages and provides indirect evidence bearing on potential donor–recipient relationships. *M. alterniflorum* exhibited the lowest mean paralog identity (94.9%), followed by *V. anagallis-aquatica* (97.2%), whereas *P. neolycopersici* displayed the highest intra-paralog identity (98.4%). Assuming broadly comparable substitution rates and evolutionary constraints, this pattern is consistent with more recent amplification of *Gnawa* in the fungal genome and is therefore more compatible with *P. neolycopersici* acting as a recipient rather than a donor. Both plant species show signatures of older activity relative to *P. neolycopersici*, with evidence for very recent activity detected in *V. anagallis-aquatica* but not in *M. alterniflorum*. Because recent or ongoing activity in a potential donor would increase the likelihood of transferring an intact, functional copy, these observations are compatible with a Lamiales-associated source, with *V. anagallis-aquatica* representing the closest sampled candidate. This interpretation is further supported by the widespread distribution of *Gnawa* homologues across the Lamiales (Fig. 5a).

Finally, phylogenetic analysis of the *Gnawa* transposase suggests that this element may have a deeper evolutionary origin beyond angiosperms (Additional file 2: Fig. S22). One plausible scenario is that an ancestral *pogo*-like element circulated in metazoan and/or fungal reservoirs before colonizing plants, eventually contributing to the more recent plant-to-fungus transfer inferred for *P. neolycopersici*. While current sampling supports this multi-step history, it does not resolve the earliest stages of *Gnawa* evolution. Broader taxonomic coverage across metazoans, fungi, and plants will be essential to distinguish among alternative scenarios.

### *Gnawa* spawned a high-copy MITE family

We identified a short, non-autonomous derivative of the *Gnawa* element (177-bp) in the genome of *P. neolycopersici*, consistent with being a Miniature Inverted-repeat Transposable Elements (MITE). This element, hereafter referred to as gMITE (*Gnawa*-derived MITE), retains the terminal inverted repeats (TIRs) but lacks the transposase-coding region. As shown in Fig. 6, gMITE is composed of a 73-bp fragment derived from the 5′ end of *Gnawa* (including the 5′ TIR) and a 104-bp fragment derived from the 3′ end (including the 3′ TIR). gMITE shares up to 99.4% nucleotide identity with the autonomous *Gnawa* element, consistent with a very recent origin. Strikingly, gMITE has undergone massive amplification within *P. neolycopersici*, reaching 172 copies in the reference genome (Additional file 1: Table S6). The expansion of gMITE was also supported by read-mapping analyses of raw, unassembled whole-genome data (PacBio and Illumina) from the two sequenced *P. neolycopersici* genomes, which revealed a marked enrichment of coverage over the 5′ and 3′ terminal regions when reads were mapped to the full-length autonomous element (Additional file 2: Fig. S23). To further investigate whether gMITE originated through a single or multiple birth events, we inferred a phylogeny using the regions shared between gMITE and autonomous *Gnawa* copies (Fig. 6). In this analysis, all gMITE sequences formed a single clade that was most closely associated with one of the nine full-length autonomous *Gnawa* copies, consistent with a single MITE birth event from an autonomous progenitor followed by a recent burst of transposition in *P. neolycopersici*, rather than multiple independent origins.

**Fig 6.**
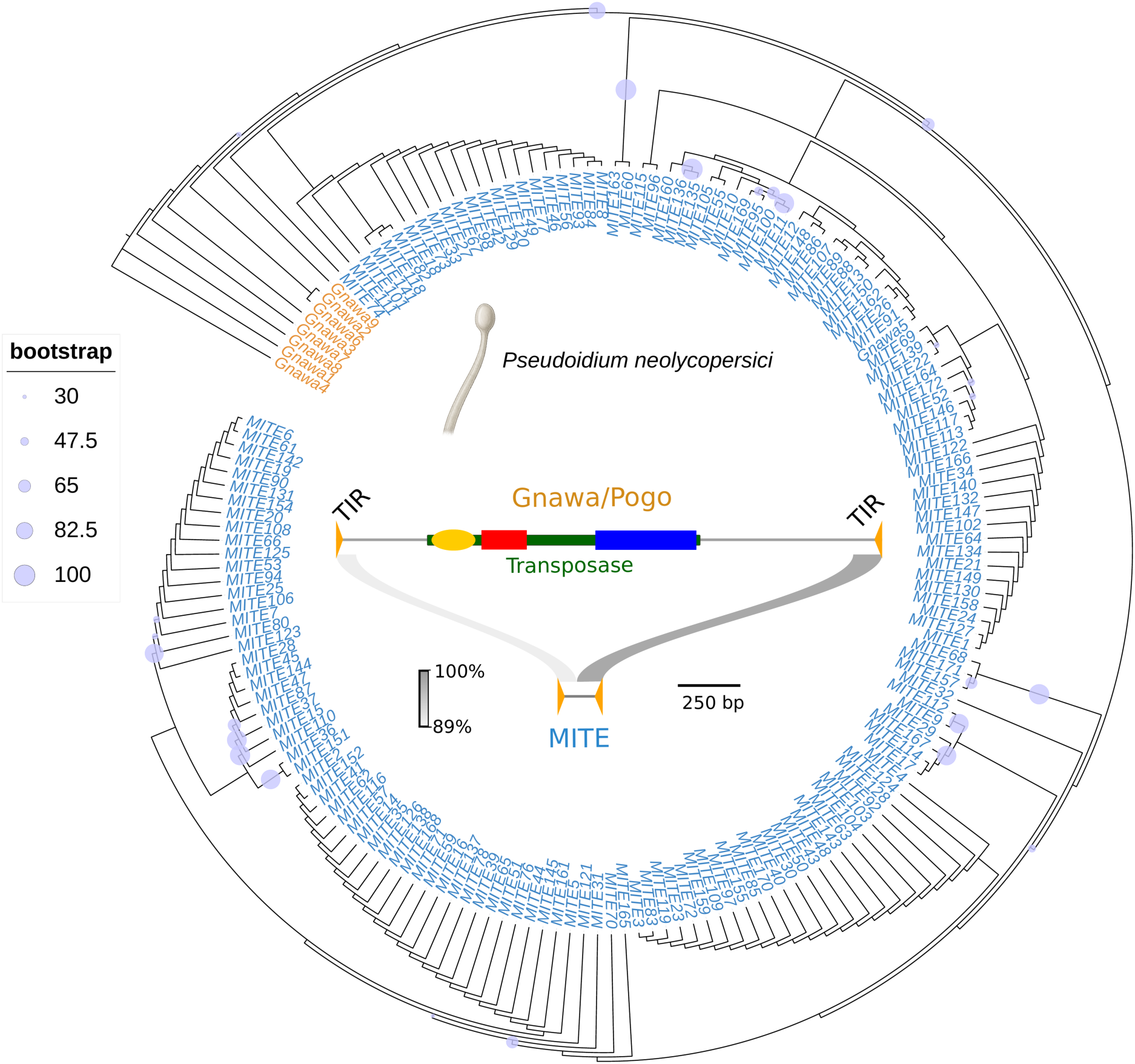
Post-transfer birth and massive amplification of a *Gnawa*-derived MITE (gMITE) in *Pseudoidium neolycopersici*. The central schematic represents the structure of the full-length autonomous *Gnawa* element (2.2-kb; domain coloring as in Fig. 4), which contains the transposase and terminal inverted repeats (TIRs), with the non-autonomous gMITE (∼177 bp) formed from its extremities. The surrounding circular phylogenetic tree displays the relationship between the full-length autonomous *Gnawa* copies and the massively amplified gMITE family (over 170 copies) within the fungal genome.

To determine whether gMITE was co-transferred from plants or instead arose *de novo* in the fungal genome after HT, we screened the two closest sampled plant species, *V. anagallis-aquatica* (the putative donor candidate) and *M. alterniflorum*, for gMITE-like sequences. However, we detected no gMITE-like sequences in either plant genome. This absence supports the hypothesis that the gMITE family originated within the fungal genome after the plant-to-fungus transfer, rather than being co-transferred from the donor lineage. To further investigate this scenario we also surveyed other genomes harbouring *Gnawa* homologues to detect *Gnawa*-derived MITEs. Among the 57 plant species containing *Gnawa*-like elements, we detected *Gnawa*-derived MITEs in only four plant species: *Geum urbanum* (370 copies), *Primula scotica* (327 copies), *Senecio aethnensis* (31 copies) and *Senecio sylvaticus* (187 copies) (Additional file 1: Table S6). Comparative alignments of these MITEs revealed differences in internal structure among lineages (Additional file 2: Fig. S27). Although the two *Senecio* species shared a similar MITE architecture consistent with a single origin in their common ancestor, MITE internal structure differed among plant species and from the gMITE identified in *P. neolycopersici*. Additionally, a phylogenetic analysis including all MITE paralogs from these plant species together with the autonomous *Gnawa* copies from *P. neolycopersici* showed that all gMITE sequences cluster specifically with the fungal *Gnawa* lineage, supporting a fungal origin of gMITE following the horizontal acquisition of the autonomous element, and arguing against an independent horizontal introduction of a pre-existing plant MITEs into *P. neolycopersici* (Additional file 2: Fig. S28).

### gMITE insertions are gene-proximal, like endogenous DNA transposons

MITEs are known to preferentially accumulate in gene-proximal regions, where they can potentially influence gene regulation [19]. We therefore examined the genomic distribution of gMITE insertions relative to genes and compared it to the endogenous TE landscape in *P. neolycopersici* (Additional file 2: Fig. S29). To establish a reference set of endogenous TEs, we performed *de novo* TE annotation of the *P. neolycopersici* genome and found that Class I retrotransposons account for 74% of total annotated TE occupancy (Additional file 1: Table S7). Because element length and scaffold gene density differ by an order of magnitude among families, and both mechanically affect distance to genes, we compared each family with the distribution expected under random placement rather than with other families. Each annotated element was repositioned uniformly at random within its own scaffold, preserving its length, and its distance to the nearest gene recomputed. Of the 172 annotated gMITE copies, 166 were located on scaffolds carrying at least one annotated gene and were retained for this analysis. All TE families were strongly depleted for insertions directly overlapping a gene (observed/expected 0.06 to 0.42), consistent with selection against gene disruption. Among elements not overlapping a gene, all families nevertheless lay closer to genes than expected by chance (Additional file 2: Fig. S30d): gMITE copies had a median distance of 2,434 bp against 3,941 bp expected (ratio 0.62; permutation p < 0.001), and 43.6% lay within 2 kb of a gene against 29.1% expected (ratio 1.50; p < 0.001) (Additional file 1: Tables S8 and S9).

We next compared the genomic distribution of gMITE insertions with that of the endogenous MITE families of *P. neolycopersici* (Additional file 2: Fig. S30). We *de novo* identified 27 endogenous MITE families comprising 2,755 copies after merging of adjacent annotations, an average of 102 copies per family. These families are of comparable length to gMITE, 202 bp against 180 bp, and occupy the same scaffolds, so the two groups share nearly identical random expectations. They showed the same pattern as gMITE, lying closer to genes than expected by chance (median distance ratio 0.68; 40.3% within 2 kb against 28.5% expected), while being more strongly depleted for direct gene overlap (observed/expected 0.18 against 0.42 for gMITE). Across all thirteen annotated TE categories, gMITE ranked fifth for gene proximity, between long interspersed nuclear elements (LINE) and Mutator-like elements (MULE-MuDR) (Additional file 2: Fig. S30d; Additional file 1: Table S8). These results indicate that gMITE is significantly gene-proximal but does not differ from the resident transposable elements of the host genome, consistent with gMITE behaving as a canonical DNA transposon following its recent birth from the horizontally acquired *Gnawa* element.

## Discussion

By screening whole genomes of 418 plants against 1,081 fungi without relying on prior gene annotation, and by combining stringent similarity filters with phylogenetic reconstruction and manual inspection of genomic context, we identified 22 high-confidence inter-kingdom HT events. While some of the detected HTs have been described previously [8, 11–13, 15], our analysis also reveals multiple additional events that went unnoticed by earlier studies because the corresponding genes may have not yet been annotated, highlighting the power of whole-genome, annotation-independent HT detection (Fig. 1). Importantly, this strategy also enabled us to document a first clear example of a recent horizontal transposon transfer from plant-to-fungus.

At the scale of the plant phylogeny, our findings reinforce the uneven and apparently rare distribution of fungus-to-plant HT across the green lineage [15], even among angiosperms, despite the availability of hundreds of high-quality genomes. However, this pattern warrants cautious interpretation. The HT events we detect likely represent only a subset of recent transfers that have occurred in the species surveyed. Rather than capturing the full scope of such events, our observations probably reflect those transfers that remain detectable in contemporary reference genomes (often derived from single individuals) because they persisted long enough in populations, potentially due to functional relevance [13] or because they have not yet been entirely lost. Supporting this perspective, several of the HTs identified here appear to be fragments of larger DNA segments that were initially integrated but subsequently underwent rapid erosion (Fig. 3c). This dynamic may also explain the apparent scarcity of plant-to-fungus transfer events. Despite screening 1,081 fungal genomes, we identified only a single unambiguous plant-to-fungus HT event and no plant-to-fungus gene transfer. Whether this reflects a genuine biological asymmetry, such as intrinsic barriers to the transfer or integration of plant DNA into fungal genomes, or simply the rapid post-transfer loss and divergence that limits detectability, remains an open question.

In the Pooideae subfamily, we found that several genes acquired from *Epichloë* through HT have undergone tandem duplications (Fig. 2b). These duplications carry two significant evolutionary implications. First, they likely reduce the risk of complete gene loss by providing functional redundancy. Second, duplicated copies may serve as raw material for subfunctionalization, at least at the expression level, as evidenced by clear differences in paralog expression patterns (Fig. 2e). By contrast, in non-vascular plants, such as *M. polymorpha*, the near absence of post-transfer duplication of fungal-derived genes (Fig. 3a) could contribute to higher long-term loss rates. This aligns with the pronounced presence– absence polymorphisms we observe across *Marchantia* subspecies and wild accessions (Additional file 2: Fig. S21). In *L. perenne*, the variation detected so far primarily involves copy-number differences between the two reference genotypes, though broader sampling will be necessary to fully assess turnover dynamics in natural populations [20]. Notably, the only HT inferred to predate the diversification of the Pooideae, a hypothetical protein locus conserved across the entire subfamily, is also the locus that has undergone the most extensive expansion across Pooideae genomes, suggesting that gene duplication contributes to the long-term retention of horizontally transferred genes in plants. Interestingly, the presence of this transferred gene in the common ancestor of the Pooideae indicates that the association between *Epichloë* fungi and Pooideae grasses predates the diversification of the subfamily, providing genomic evidence that this symbiosis is at least ∼20 million years old [21, 22]. More broadly, our results illustrate how HT can preserve a genomic footprint of ancient ecological interactions, providing a powerful means to infer the evolutionary age of symbiotic associations.

It is noteworthy that the majority of the transferred coding genes identified in this study are predicted to encode enzymes or enzyme-associated domains, including glycoside hydrolases, glutathione S-transferases, acetyltransferases, phosphatases, and FAD-dependent monooxygenase modules. This pattern aligns with previous observations that horizontally acquired genes in plants frequently encode enzymes, particularly glycoside hydrolases [10, 12, 23]. While this apparent bias may reflect a biologically meaningful trend, it must be interpreted with caution. One plausible explanation is that enzymatic genes are more likely to be retained post-transfer because they often provide discrete biochemical functions that require minimal integration into lineage-specific interaction networks. Unlike components of tightly co-evolved multi-protein complexes or regulatory pathways, enzymes can more easily be “plugged into” existing cellular physiology, offering an immediate selective advantage. This interpretation resonates with the “complexity hypothesis”, initially proposed based on prokaryotic HT analyses [24] and later supported by empirical evidence showing that highly connected proteins are less frequently transferred [25]. In this context, enzymes involved in detoxification or specialized metabolism may be particularly prone to retention if they confer context-dependent fitness benefits, as documented for horizontally acquired metabolic and stress-related genes in eukaryotes [6, 26]. However, the molecular functions and adaptive significance of these fungal-derived genes in both bryophytes and Pooideae remain to be experimentally validated through functional assays.

Our results also indicate that these transfers were probably DNA-mediated, and included not only the coding sequences but also the genomic context of the transferred genes, which may favor their transcriptional reactivation. In Pooideae, we detect cases in which transferred sequences extend beyond coding boundaries and include introns. At the same time, our analysis also points to rapid post-transfer remodeling in the recipient genome. For the glycoside hydrolase paralogs in *L. perenne* for instance, at least two independent intron replacement events occurred after duplication, including one case involving an endogenous MITE-derived insertion, illustrating how transferred gene structures can be rapidly reshaped. Such remodeling may facilitate accommodation of foreign genes into new genomic environments by altering splicing architecture and gene regulation, while the extensive erosion of flanking regions in older transfers underscores that non-coding sequences may be lost quickly unless constrained.

The consistently low expression of some HT-derived loci, particularly those encoding the hypothetical protein family that has undergone extensive duplication in Pooideae and has been maintained over long evolutionary timescales, is intriguing. Low basal or absence of expression does not necessarily imply lack of function, as expression may be restricted to specific tissues, developmental stages, or environmental conditions that are not represented in the available datasets. More generally, HT-derived genes may exhibit conditional activation during biotic interactions in Pooideae. For example, the *Epichloë*-derived gene *Fhb7* in *Thinopyrum* has been reported to be induced in response to *Fusarium* infection [13].

Finally, this study provides evidence for a horizontal transfer event in which a plant TE crossed kingdom boundaries and integrated into the genome of the obligate biotroph *P. neolycopersici*. Following its acquisition, *Gnawa* has been reactivated and extensively amplified, such that the fungus now harbours about 180 copies of plant-derived sequences. Notably, no homologues of the transferred *Gnawa* element were detected in tomato (*Solanum lycopersicum*), the primary present-day host of *P. neolycopersici,* nor in closely related *Solanum* species. This observation suggests that the acquisition event did not involve the fungus’s currently main host, but instead may have occurred in a broader ecological context, extending beyond the canonical *P. neolycopersici*–tomato interaction. In nature, *P. neolycopersici* is able to infect a wide range of plant species beyond tomato, including representatives of several plant families and orders such as Lamiales and Cucurbitales, among others [27]. This broad host range may have increased the opportunities for cross-species DNA transfer of *Gnawa* between the fungus and diverse plant genomes. The unusually high sequence identity observed between fungal copies of *Gnawa* and homologues from multiple, taxonomically distant plant lineages is intriguing and could suggest the occurrence of multiple horizontal transfer events. However, it remains unclear whether *P. neolycopersici* is best viewed as a terminal recipient in this putative circulation network, or whether it might contribute to further redistribution of this element playing the role of a vector. Discriminating between these scenarios and others will require denser taxonomic sampling across candidate donor and recipient lineages. Nonetheless, our findings highlight powdery mildew fungi, and *Gnawa* in particular, as promising model systems for investigating the ecological and mechanistic conditions that enable cross-kingdom TE transfer. They also raise questions about the biological impact of *Gnawa* acquisition on *P. neolycopersici*, including possible consequences for genome dynamics, adaptation, pathogenicity, and the fine-tuning of interactions with its host plants.

## Conclusion

Our comparative genomic analysis reveals a sparse, asymmetric, and phylogenetically uneven landscape of recent horizontal DNA transfers between plants and fungi, shaped by distinct ecological associations, with *Epichloë* and the *Pleosporales* emerging as recurrent donors for grasses and non-vascular plants, respectively. Importantly, our genome-wide approach shows that horizontal transfer is not limited to genes but can involve larger genomic regions that are subsequently eroded and reshaped, while conserved genes may be duplicated, diverge, or be lost in their new hosts.

The plant-derived *Gnawa* transposon illustrates an even more dynamic outcome: after being transferred to *Pseudoidium neolycopersici*, it remained active and gave rise de novo to a family of MITEs that has recently undergone a transpositional burst. Thus, the evolutionary history of horizontally transferred DNA does not end with its acquisition. Once introduced into a new genome, the foreign DNA embarks on a new evolutionary trajectory in which conservation, erosion, remodeling, duplication, loss, and amplification determine its ultimate fate and its contribution to genomic diversification.

## Methods

### Genome acquisition and database construction

To perform a comprehensive bidirectional survey of HT, we compiled a genome dataset comprising 1,081 fungal genomes retrieved from JGI MycoCosm [28] and NCBI [29], together with 418 plant genomes obtained from Phytozome [30] and NCBI (Additional file 1: Tables S1 and S2). The dataset was curated to maximize taxonomic coverage and included representatives of the major fungal lineages (Ascomycota, Basidiomycota, and early-diverging fungi) and plant lineages, including angiosperms, gymnosperms, lycophytes, monilophytes, bryophytes, chlorophytes, charophytes, and rhodophytes.

### Genome-wide homology search and primary filtering

All-vs-all pairwise comparison was performed by aligning fungal genomes against plant genomes using BLASTn (BLAST+ suite) [31]. Only High-scoring Segment Pairs (HSPs) exhibiting a nucleotide sequence identity ≥ 70% and an alignment length ≥ 500 bp were retained. Alignments were only retained if they were located on scaffolds larger than 20 kbp in both the query (fungal) and subject (plant) genomes.

### Filtering of tandem repeats, ribosomal and organellar DNA

Candidate sequences were extracted based on the filtered BLAST coordinates. To remove low-complexity regions, we processed the sequences using Tandem Repeats Finder (TRF) [32]. This approach minimizes the retention of spurious similarity driven by microsatellites or simple sequence repeats, which are common in eukaryotic genomes. Candidate sequences were further filtered by screening against nucleotide databases of ribosomal RNA genes (SILVA) [33] and organellar genomes (plastid and mitochondrial sequences, NCBI). BLASTn searches were performed, and candidates showing significant similarity to ribosomal or organellar DNA were removed. This step ensures that only nuclear genomic regions are retained, as ribosomal and organellar sequences are known to produce strong cross-kingdom similarity signals unrelated to horizontal transfer.

### Clustering and functional annotation

To reduce redundancy among the detected hits, candidate sequences were clustered using CD-HIT-EST [34] with a sequence identity threshold of 80% (-c 0.8), retaining the longest sequence as the representative of each cluster. Representative sequences were functionally annotated using EggNOG-mapper in DIAMOND mode [35]. To minimize false-positive HT predictions, candidates annotated as highly conserved housekeeping genes were excluded using a keyword-based filter applied to the EggNOG annotations. This filter removed sequences associated with ubiquitous cellular functions, including ribosomal proteins, ATPases, actins, tubulins, polymerases, ubiquitins, GAPDH, HSP70, PRP8, and related proteins. These genes are broadly conserved across eukaryotes and frequently produce spurious cross-kingdom sequence similarities that can be mistaken for HT. Although this conservative filtering strategy may discard a small number of genuine HT events, it substantially reduces the number of false positives and decreases the computational burden of subsequent homology searches, phylogenetic reconstruction, and manual curation.

### Homolog retrieval

For each remaining candidate, homologous sequences were retrieved from both plant and fungal genomes using BLASTn (BLAST+ suite) [31]. Homologs were required to meet a minimum nucleotide identity of 70% and a minimum query coverage of 60%. A maximum of top five best hit homologous sequences per genome was retained to limit redundancy. Only homologs located on scaffolds of at least 20 kb were kept. Homologous sequences were grouped into multi-FASTA files, one per candidate locus.

### Contaminated genome removal

To identify potential assembly contaminants, the taxonomic composition of the homologue sets was examined across all candidate loci. Plant or fungal genomes that repeatedly appeared as the sole or predominant source of cross-kingdom homologues in a large number of independent candidate datasets were flagged as potential contaminants. For each flagged genome, the genomic distribution of candidate loci was then inspected. Candidate loci distributed across multiple independent scaffolds were interpreted as evidence of genome-wide contamination, leading to exclusion of the corresponding species from all homologue datasets. In contrast, when candidate loci consistently mapped to a single scaffold or chromosome, only that scaffold or chromosome was excluded. Several removed chromosomes corresponded to unanchored chromosomes (ChrUn). Overall, this iterative curation identified 15 contaminated species or genomic scaffolds and reduced the number of HT candidate loci by 98.99% before phylogenetic analysis.

### Enrichment, multiple alignment and phylogenetic analysis

To increase taxonomic sampling and phylogenetic resolution, an enrichment step was performed using the NCBI core_nt nucleotide database. Taxonomic identifiers corresponding to plant and fungal species (Taxonomic Database NCBI) were used to get fungi and plant homologs using BLASTn (BLAST+ suite) [31]. This step improves robustness of candidate evaluation by incorporating additional homologous sequences. A multi-representative querying strategy was used: for each cluster, up to five representative sequences per kingdom were selected, one per species, choosing the longest sequence within each species and ranking representatives by length descending, and independently queried against core_nt using BLASTn (e-value 1e-50, identity ≥ 70%, query coverage ≥ 60%). To ensure balanced phylogenetic representation, hits were strictly filtered by kingdom, capping the maximum number of hits per species at 5 and the total number of sequences per kingdom at 150. These sequences were merged with plants and fungi homologs resulting from whole-genome comparisons. For each cluster, Multiple Sequence Alignments (MSA) were generated using MAFFT v7.490 (--auto) [36] and cleaned with trimAl v1.4 (-automated1, keeping headers) [37]. Maximum Likelihood trees were inferred using IQ-TREE 2 [38] with 1,000 Ultrafast Bootstrap replicates (-bb 1000) [39], with the substitution model automatically selected by ModelFinder [40].

### Topology analysis

Phylogenetic trees were midpoint-rooted and manually evaluated for evidence of incongruence between plant and fungal lineages. Candidates were considered consistent with HT when they displayed either (i) a well-supported mixed plant–fungus clade (bootstrap support ≥70%) or (ii) a restricted taxonomic distribution in one kingdom associated with a well-supported placement among homologues from multiple species of the other kingdom. The evolutionary breadth of the sampled taxa was also considered: homologues shared by several distantly related species of the same kingdom and forming a phylogenetically coherent group were considered more consistent with ancient vertical inheritance than with HT. Trees showing weakly supported or unresolved relationships, or insufficient taxonomic representation, were excluded.

### Annotation of validated horizontal transfer loci

Validated transferred loci were annotated by predicting putative coding sequences directly from the transferred genomic segments. Each sequence was translated in all six reading frames using EMBOSS getorf [41] to identify open reading frames (minimum length of 150 bp) and their corresponding protein sequences. Predicted proteins were functionally annotated using InterProScan (v5.74-105.0) [42] to identify conserved protein domains. The resulting annotations were used to reconstruct domain architectures and generate schematic representations of the transferred genes. For candidate proteins lacking InterPro annotations, conserved domains were further investigated using the NCBI Conserved Domain Search (CD-Search) web server [43] with default parameters.

### RNA-seq read mapping and gene expression quantification

Publicly available paired-end RNA-seq datasets were downloaded from the NCBI Sequence Read Archive (SRA) (Additional file 1: Table S4). Adapter sequences and low-quality bases were removed using Trimmomatic [44]. Filtered reads were aligned to the corresponding reference genomes using HISAT2 (v2.2.1) [45, 46] with default parameters. Gene-level read counts were generated using featureCounts (v2.0.1) [47] based on the corresponding genome annotations. Transcript abundance was normalized as Transcripts Per Million (TPM) to account for differences in gene length and sequencing depth. TPM values were calculated only for genomes with available gene annotations and were used to assess the expression of HT-derived genes across the analyzed RNA-seq datasets.

### *Pogo*/*Gnawa* phylogeny

*Pogo*/*Gnawa* transposase sequences were predicted by translating genomic copies of *Gnawa* homologs and paralogs identified in all analyzed plant genomes and in *Pseudoidium neolycopersici*. These sequences were combined with the complete set of *Tc1*/*mariner* transposases previously identified from viridiplantae, metazoa, and fungi by Dupeyron et al. [18]. Protein sequences were aligned using MAFFT v7.505 with default parameters. Poorly aligned positions were removed using trimAl v1.5.rev0 [37] with default settings. Maximum-likelihood phylogenetic inference was performed using IQ-TREE 2 [38] with automatic model selection (-m MFP) and branch support was assessed using 1,000 ultrafast bootstrap replicates (-bb 1000).

### Variation of horizontal transfer among wild *Marchantia* accessions

To investigate intraspecific variation of HT-derived sequences, whole-genome sequencing data from nine *Marchantia* accessions, representing three accessions from each of the three recognized subspecies and spanning contrasting geographic origins (Additional file 1: Table S5), were analyzed. Paired-end genomic reads were mapped to the *M. polymorpha* reference genome using Minimap2 (v2.26) [48] with the short-read preset (-ax sr). Read alignments were subsequently inspected to validate the presence/absence patterns inferred from the genome assemblies and to assess the sequence conservation of HT-derived loci across the nine accessions.

### Transposable element annotation of *Pseudoidium neolycopersici*

Transposable elements (TEs) in the *Pseudoidium neolycopersici* genome were annotated using a *de novo* TE detection approach. First, a *de novo* repeat library was generated using RepeatModeler2 [49]. The resulting consensus sequences were subsequently refined and curated using TEtrimmer [50], which extends fragmented consensus sequences, removes redundancy, and improves TE family classification through iterative sequence extension, multiple-sequence alignment, TE boundary definition, and consensus reconstruction. Curated TE consensus sequences were classified based on their structural characteristics and sequence similarity to known TE families. Miniature inverted-repeat transposable elements (MITEs) were identified using MITE-Tracker [51], which detects MITE families based on their characteristic structural features, including terminal inverted repeats (TIRs) and target site duplications (TSDs). The resulting MITE consensus sequences were manually inspected to remove redundant and low-confidence predictions, then combined with the TEtrimmer-curated consensus sequences into a single non-redundant library. This library was used as a custom database for genome-wide annotation with RepeatMasker [52], enabling the identification and classification of all genomic copies (paralogs) belonging to each TE family, including both autonomous and non-autonomous elements. Because a single element is frequently reported as several adjacent RepeatMasker hits, annotations assigned to the same family and separated by less than 500 bp were merged into single elements using bedtools merge (v2.31.1) [53].

### Analysis of MITE distribution relative to genes

The distance from each merged element to the nearest annotated gene was computed with bedtools closest (v2.31.1). Where a single element was reported against several equidistant genes, one observation was retained per element. Elements located on scaffolds carrying no annotated gene were excluded. Insertion preferences were assessed against a random-placement null: each element was repositioned uniformly at random within its own scaffold, preserving its length, and its distance to the nearest gene recomputed; 300 to 3,000 permutations were performed per family according to copy number. Two-sided empirical p values were derived from the permutation distributions of the proportion of elements overlapping a gene and, among non-overlapping elements, of the median distance and of the proportion lying within 2 kb of a gene. MITE families were treated as a separate group and were not included in the Class II group.

The nucleotide sequences of the horizontally transferred loci and of the *Gnawa* copies identified in *P. neolycopersici* are deposited in figshare [54] and are also provided as Additional file 3. The complete detection workflow described above, from the pairwise genome comparisons to the phylogenetic evaluation of the candidates, is implemented in the BRIDGE (Bidirectional Recognition and Identification of Genomic Exchanges) pipeline; the version used for the analyses reported here (v1.0.0) is archived in Zenodo [55].

## Supporting information

Additional file 2. Supplementary Figures S1-S30 with legends

Additional file 3. FASTA sequences of the horizontally transferred loci and of the Gnawa copies

## Declarations

### Ethics approval and consent to participate

Not applicable.

### Consent for publication

Not applicable.

### Availability of data and materials

The datasets supporting the conclusions of this article are included within the article and its additional files. The genome assemblies analysed are publicly available from NCBI, Phytozome and the JGI MycoCosm portal and are listed in Additional file 1 (Tables S1 and S2); the *P. neolycopersici* MF1 v1.0 assembly is MycoCosm portal Psene1. The RNA-seq and whole-genome sequencing runs analysed are listed in Additional file 1 (Tables S4 and S5), and the raw reads used to validate the integration of *Gnawa* in *P. neolycopersici* are available in the NCBI Sequence Read Archive under accessions SRR11557659, SRR11557660, SRR11557661, SRR11557663, SRR11557664 and SRR4017876. The FASTA sequences of the horizontally transferred loci and of the *Gnawa* copies identified in this study are available in the figshare repository [54] and in Additional file 3.

The BRIDGE (Bidirectional Recognition and Identification of Genomic Exchanges) pipeline developed in this study is publicly available on GitHub [14]. The version used for the analyses reported in this study (v1.0.0) has been permanently archived on Zenodo [55].

- Project name: BRIDGE-HT-Pipeline
- Project home page: https://github.com/Lee-06/BRIDGE-HT-Pipeline
- Archived version: https://doi.org/10.5281/zenodo.22641750
- Operating system(s): Linux (Ubuntu 22.04)
- Programming language: Python, Bash
- Other requirements: BLAST+, samtools, Tandem Repeats Finder, CD-HIT-EST, eggNOG-mapper, TaxonKit, MAFFT, trimAl, IQ-TREE 2; Python 3.7 or later with pandas and Biopython
- License: MIT
- Any restrictions to use by non-academics: none

### Authors’ contributions

ME conceived the project. LM performed the analyses and developed the HT detection pipeline with methodological input from ME. EL performed gene annotation. JQ, SK, and RP performed TE annotation in *P. neolycopersici* and contributed to the interpretation of the results. JHH and KDK performed domain analyses and classified the *Gnawa* element. AG contributed to data interpretation. LM and ME wrote the manuscript with input from all authors. All authors read and approved the final manuscript.

## Acknowledgements

We thank the U.S. Department of Energy Joint Genome Institute (JGI) and the principal investigators of the MycoCosm and Phytozome genome projects whose data were analysed in this study. The work conducted by the U.S. Department of Energy Joint Genome Institute, a DOE Office of Science User Facility, is supported by the Office of Science of the U.S. Department of Energy operated under Contract No. DE-AC02-05CH11231.

## Supplementary tables (Additional File 1)

**Supplementary Table 1. Plant species used in the genome-wide screen.** Taxonomic assignment of the plant genome assemblies retrieved from NCBI and Phytozome and included in the pairwise fungal–plant comparisons. For each species, the table provides the species name and its taxonomic classification.

**Supplementary Table 2. Fungal genome assemblies used in the genome-wide screen.** The 1,081 fungal genome assemblies retrieved from JGI MycoCosm and NCBI. For each assembly the table gives the portal assembly code, the species name, whether a taxonomic match was recovered, the rank at which the match was made, and the complete resolved lineage.

**Supplementary Table 3. Inventory of the horizontal transfer events identified in this study.** For each event, the table lists the event identifier, the corresponding Supplementary Figure, the abbreviation and colour code used in Figs. 2 and 3, the predicted function, the recipient gene identifier (when available), the recipient species or taxonomic group, the inferred putative fungal donor and its taxonomic assignment, whether the event has been reported previously, and the corresponding reference DOI.

**Supplementary Table 4. Publicly available RNA-seq datasets used for expression quantification.** The 59 RNA-seq SRA runs analysed across 17 species, spanning different tissues and stress conditions. For each run the table gives the SRA run accession, the species, the tissue or experimental condition, the library layout, and the total number of reads.

**Supplementary Table 5. *Marchantia polymorpha* wild accessions used to validate presence and absence polymorphism.** The nine whole-genome sequencing datasets analysed, comprising three accessions for each of the three recognised subspecies (*montivagans*, *polymorpha* and *ruderalis*) sampled from contrasting geographic origins. For each dataset the table gives the SRA run accession, the species, the subspecies, the accession name, the geographic zone of origin, and the number of reads.

**Supplementary Table 6. Genomic coordinates of *Gnawa* elements and *Gnawa*-derived MITEs.** Positions of the 1,096 annotated copies, comprising the nine full-length autonomous *Gnawa* elements and the 172 *gMITE* copies identified in *Pseudoidium neolycopersici*, together with the *Gnawa*-derived MITEs detected in four plant species (*Geum urbanum*, 370 copies; *Primula scotica*, 327; *Senecio sylvaticus*, 187; *Senecio aethnensis*, 31). For each copy the table gives the copy identifier, the scaffold or chromosome, the start and end coordinates, the assembly used, and the species.

**Supplementary Table 7. Transposable element content of the *Pseudoidium neolycopersici* genome.** Genomic occupancy of each transposable element superfamily recovered by the de novo annotation, grouped by class, given in base pairs and as a percentage of total annotated TE occupancy.

Supplementary Table 8. Insertion preferences of transposable element families in *Pseudoidium neolycopersici* relative to random expectation. Comparison of the observed genomic position of each annotated element with the distribution expected under random placement, obtained by repositioning every element uniformly at random within its own scaffold while preserving its length. This null controls for element length and for scaffold gene density, which differ by an order of magnitude among families. Two aspects are reported separately: the proportion of elements directly overlapping a gene, and, among elements that do not overlap a gene, the median distance to the nearest gene together with the proportion lying within 2 kb of a gene. For each family the table gives the number of copies analysed, the median element length, and, for each of the three statistics, the observed value, the mean value expected under permutation, their ratio, and the two-sided empirical *P* value. The number of permutations is given in the last column; the smallest attainable *P* value is 2/(*B* + 1). Elements located on scaffolds carrying no annotated gene were excluded, and a single observation was retained per element. MITE families are reported separately and are not included in the Class II group.

**Supplementary Table 9. Distance to the nearest gene for each transposable element category in *Pseudoidium neolycopersici*.** Copy number and summary statistics of the distance separating each annotated element from the nearest gene, given in base pairs, for the eleven element superfamilies recovered by the de novo annotation, together with *gMITE* and with the endogenous MITE families considered as a single group. For each category the table gives the number of copies, the mean, median, standard deviation, minimum, first and third quartiles, and maximum distance. Categories are mutually exclusive; the MITE row comprises the endogenous MITE families, which are reported separately from the Class II superfamilies.

## Supplementary figures (Additional File 2)

**Supplementary Figure 1. Maximum-likelihood phylogeny of the FAD-binding monooxygenase (FMO)**. The phylogeny was inferred from nucleotide sequences. Grey triangles represent collapsed fungal clades, with labels indicating the number of represented species. HT-derived FMO copies from *Lolium perenne* are shown in green. Circle size is proportional to ultrafast bootstrap support.

**Supplementary Figure 2. Maximum-likelihood phylogeny of the hypothetical protein (*HP*).** The phylogeny was inferred from nucleotide sequences. HT-derived HP gene copies from the Pooideae species are shown in green. Circle size is proportional to ultrafast bootstrap support. HGT1 denotes the ancestral acquisition in the common ancestor of the Pooideae; HGT2 and HGT3 denote the two independent, more recent transfers.

**Supplementary Figure 3. Maximum-likelihood phylogeny of the glycoside hydrolase (*GH*) gene**. The phylogeny was inferred from nucleotide sequences. Grey triangles represent collapsed fungal clades, with labels indicating the number of represented species. HT-derived GH copies from the Pooideae species are shown in green. Circle size is proportional to ultrafast bootstrap support.

**Supplementary Figure 4. Maximum-likelihood phylogeny of the Oxopyrrolidines (*OP*) gene**. The phylogeny was inferred from nucleotide sequences. *Lolium perenne* homolog is shown in green. Circle size is proportional to ultrafast bootstrap support.

**Supplementary Figure 5. Maximum-likelihood phylogeny of the glutathione S-transferase (*Fhb7*) gene**. The phylogeny was inferred from nucleotide sequences. *Thinopyrum elongatum* homolog is shown in green. Circle size is proportional to ultrafast bootstrap support.

**Supplementary Figure 6. Maximum-likelihood phylogeny of the protein phosphatase (*PP*) gene**. The phylogeny was inferred from nucleotide sequences. *Marchantia polymorpha* homologs are shown in green. Grey triangles represent collapsed fungal clades, with labels indicating the number of represented species. Circle size is proportional to ultrafast bootstrap support.

**Supplementary Figure 7. Maximum-likelihood phylogeny of a hypothetical protein (*HP*) gene.** The phylogeny was inferred from nucleotide sequences. *Marchantia polymorpha* homologs are shown in green. Grey triangles represent collapsed fungal clades, with labels indicating the number of represented species. Circle size is proportional to ultrafast bootstrap support.

**Supplementary Figure 8. Maximum-likelihood phylogeny of PITH & SM-like domain (*PITH-SM*) gene.** The phylogeny was inferred from nucleotide sequences. *Marchantia polymorpha* homologs are shown in green. Grey triangles represent collapsed fungal clades, with labels indicating the number of represented species. Circle size is proportional to ultrafast bootstrap support.

**Supplementary Figure 9. Maximum-likelihood phylogeny of phenolic acid decarboxylase and FAD-dependent monooxygenase (*PAD-FMO*) gene. The phylogeny was inferred from nucleotide sequences.** *Marchantia polymorpha* homolog is shown in green. Grey triangles represent collapsed fungal clades, with labels indicating the number of represented species. Circle size is proportional to ultrafast bootstrap support.

**Supplementary Figure 10. Maximum-likelihood phylogeny of a hypothetical protein (*HP*) gene.** The phylogeny was inferred from nucleotide sequences. *Marchantia polymorpha* and *Ceratodon purpureus* homologs are shown in green. Circle size is proportional to ultrafast bootstrap support.

**Supplementary Figure 11. Maximum-likelihood phylogeny of glutathione S-transferase (*GST*) gene.** The phylogeny was inferred from nucleotide sequences. Plant homologs are shown in green. HGT1–HGT5 indicate five independent HT events involving *GST* genes. Circle size is proportional to ultrafast bootstrap support.

**Supplementary Figure 12. Maximum-likelihood phylogeny of the acetyltransferase (*AT*) gene.** The phylogeny was inferred from nucleotide sequences. Plant homologs are shown in green. HGT1 and HGT2 indicate two independent HT events of the *AT* gene. Circle size is proportional to ultrafast bootstrap support.

**Supplementary Figure 13. Maximum likelihood phylogenetic tree of *Gnawa* element displaying branch lengths.** The phylogeny was inferred from nucleotide sequences. The unusually short branch lengths observed between *P. neolycopersici* sequences and distantly related eudicot lineages highlight the exceptionally high nucleotide identity maintained across these elements.

**Supplementary Figure 14. InterProScan annotation of transferred genes containing conserved domains.** Transferred DNA segments were annotated for coding sequences (CDSs) and conserved domains as described in the Materials and Methods section. In each panel, the top line represents the transferred DNA segment (in nucleotides). Grey arrows indicate predicted coding sequences (CDS_nn) containing conserved domains, and coloured boxes show the positions of the detected domains within each CDS. Abbreviations: Lp, *Lolium perenne*; Mp, *Marchantia polymorpha*; Te, *Thinopyrum elongatum*; Pp, *Physcomitrium patens*; Ms, *Monoraphidium* sp. 549. FMO, FAD-binding monooxygenase; HP, hypothetical protein; OP, oxopyrrolidines; GH, glycoside hydrolase; PP, protein phosphatase; PITH-SM, PITH & SM-like domain; PAD-FMO, phenolic acid decarboxylase/FAD-dependent monooxygenase; Fhb7, glutathione S-transferase Fhb7; AT, acetyltransferase.

**Supplementary Figure 15. Chromosomal distribution of HT-derived genes across representative Pooideae genomes.** HT-derived genes are coloured according to gene family (*FMO*, FAD-binding monooxygenase; *HP*, hypothetical protein; *GH*, glycoside hydrolase; *OP*, oxopyrrolidine; *NG*, non-genic; *Fhb7*, glutathione S-transferase). Asterisks indicate tandemly duplicated copies.

**Supplementary Figure 16. Genomic context and flanking sequence conservation of horizontally transferred genes in *Lolium perenne*.** Sequence identity plots comparing *Epichloë* donor genomes to the *L. perenne* genome for the oxopyrrolidines (*OP*) and hypothetical protein (*HP*) loci. Percentage represents Blastn sequence identity.

**Supplementary Figure 17. RNA-seq read count of the hypothetical protein (HP) and glycoside hydrolase (GH) families in the Pooideae.** Bar plots show leaf RNA-seq read counts for the corresponding HT-derived grass homologs. Species names shown in grey indicate that no leaf RNA-seq data were available. Circle size is proportional to ultrafast bootstrap support.

**Supplementary Figure 18. Intraspecific copy number variation of horizontally acquired loci in *Lolium perenne*.** Chromosomal mapping compares the presence, absence, and tandem duplication status of the HT-derived genes (*FMO*, *GH*, *HP*, *OP*, and *NG*) between two high-quality *L. perenne* reference genomes, Kyuss and P226/135/16.

**Supplementary Figure 19. Expression profiles of horizontally transferred genes in non-vascular plants.** Log₁₀-transformed RNA-seq read counts of HT-derived genes (*HP*, hypothetical protein; *AT*, acetyltransferase; *GST*, glutathione S-transferase) across diverse tissues, developmental stages, and experimental conditions in *Ceratodon purpureus*, *Lunularia cruciata*, *Physcomitrium patens*, *Syntrichia caninervis*, *Pseudopediastrum boryanum*, and *Tetradesmus obliquus*. Pie charts indicate the proportion of mapped and unmapped reads, and numbers above each chart indicate the total number of RNA-seq reads (millions).

**Supplementary Figure 20. Structural organization and transcriptional evidence of representative HT loci in *Marchantia polymorpha*. a)** Genomic organization of the PAD-FMO locus showing its conservation with the putative fungal donor (*Delphinella strobiligena*) and RNA-seq read coverage across the transferred region in *M. polymorpha*. **b)** Genomic organization of the *PITH-SM* locus showing its conservation with the putative fungal donor (*Corynespora cassiicola*) and its integration into a pre-existing endogenous host gene (MpTK2_5g06260), resulting in a chimeric locus.

**Supplementary Figure 21: Intraspecific presence/absence polymorphism of horizontal gene transfers across *Marchantia polymorpha* wild accessions.** Read coverage plots for wild accessions from the three major subspecies (*montivagans*, *polymorpha*, and *ruderalis*), confirming that differences in HT content (*HP*, *PITH*-*SM*, *GST*, *PP*, and *PAD*-*FMO*) reflect genuine biological variation and ongoing gene loss rather than assembly artifacts.

**Supplementary Figure 22: Phylogenetic placement of the *Gnawa* transposable element.** Maximum likelihood tree based on the transposase domain of known *Tc1/mariner* elements, placing the *Gnawa* transposon within the *pogo*-like clade typical of Viridiplantae (plants) and distinct from typical fungal elements.

**Supplementary Figure 23. Read coverage profiles of the *Gnawa* element across independent *Pseudoidium neolycopersici* genome sequencing datasets.** Whole-genome Illumina and PacBio sequencing reads from two independent *P. neolycopersici* isolates map continuously across the full-length *Gnawa* consensus sequence.

**Supplementary Figure 24. PacBio long reads spanning the *Gnawa* terminal inverted repeat (TIR) junctions.** Long sequencing PacBio reads cover the entire transposon and extend into the flanking fungal genomic regions, unambiguously demonstrating the physical integration of the *Gnawa* element within the *P. neolycopersici* genome.

**Supplementary Figure 25. Pairwise nucleotide sequence identity of full-length *Gnawa* homologs across different plant orders.** The heatmap demonstrates the remarkably high sequence conservation (frequently exceeding 95-99% identity) among *Gnawa* copies from the fungal recipient (*P. neolycopersici*) and diverse, distantly related plant hosts.

**Supplementary Figure 26. Intraspecies paralog divergence of *Gnawa* homologs across host plants and *P. neolycopersici*.** Density distributions of intra-genomic sequence identity illustrate minimal divergence among *Gnawa* paralogs in several lineages, indicating recent transpositional activity and lineage-specific expansion.

**Supplementary Figure 27. Comparative sequence alignment of *Gnawa*-derived MITEs from plants and fungi.** The alignment compares the internal structures of MITEs identified in *Pseudoidium neolycopersici* and in the plant species *Geum urbanum*, *Primula scotica*, and *Senecio* spp. Terminal inverted repeats (TIRs) are highly conserved, whereas the internal regions differ substantially among lineages, consistent with independent MITE formation events.

**Supplementary Figure 28. Phylogenetic relationship of *Gnawa*-derived MITEs across lineages.** The phylogenetic tree includes all MITE paralogs from plants and *P. neolycopersici*.

**Supplementary Figure 29. Distance to nearest genes for *gMITE* and endogenous transposable elements (TEs) in *P. neolycopersici*.** The boxplot compares the insertion proximity to genes for *gMITE* against the broader TE landscape, categorized by subclass into Class I retrotransposons (LTRs, non-LTRs) and other Class II DNA transposons. Element length differs by an order of magnitude among families and is controlled for in the permutation analysis (Supplementary Table 8).

**Supplementary Figure 30. Insertion landscape of *gMITE* and of the endogenous MITE families of *P. neolycopersici* relative to random expectation. a, b,** Cumulative distribution of the distance separating each element from the nearest annotated gene, for the 166 *gMITE* copies (a) and the 2,755 copies belonging to the endogenous MITE families (b). Coloured lines show the observed distribution; the grey band and dashed line show the distribution expected under random placement, obtained by repositioning every element uniformly at random within its own scaffold while preserving its length (mean and 95% range across 3,000 permutations for *gMITE* and 1,200 for the endogenous families). c, Ratio of observed to expected values for three statistics: the proportion of elements directly overlapping a gene, and, among elements that do not overlap a gene, the median distance to the nearest gene and the proportion lying within 2 kb of a gene. Grey bars give the 95% range of the permutation distribution, centred on a ratio of 1 (dashed line). d, The same two ratios for all thirteen TE categories, ordered by median-distance ratio. Filled circles denote *P* < 0.05; open circles denote non-significant deviations, all in families with fewer than 60 copies. Every family is depleted for insertions overlapping a gene and lies closer to genes than expected by chance, with *gMITE* ranking fifth of thirteen.

## Additional files

**Additional file 1** (.xlsx). Supplementary Tables S1–S9. Table S1: plant species used in the genome-wide screen. Table S2: fungal genome assemblies used in the genome-wide screen. Table S3: inventory of the horizontal transfer events identified in this study. Table S4: publicly available RNA-seq datasets used for expression quantification. Table S5: *Marchantia polymorpha* wild accessions used to validate presence and absence polymorphism. Table S6: genomic coordinates of *Gnawa* elements and *Gnawa*-derived MITEs. Table S7: transposable element content of the *Pseudoidium neolycopersici* genome. Table S8: insertion preferences of transposable element families in *P. neolycopersici* relative to random expectation. Table S9: distance to the nearest gene for each transposable element superfamily in *P. neolycopersici*.

**Additional file 2** (.pdf). Supplementary Figures S1–S30, with legends. Maximum-likelihood phylogenies of the horizontally transferred genes (Figs. S1–S12) and of *Gnawa* (Figs. S13 and S22); domain annotation of the transferred segments (Fig. S14); genomic organization, flanking-sequence conservation, expression and copy number variation of the transferred genes (Figs. S15–S20) and their presence–absence polymorphism among *M. polymorpha* accessions (Fig. S21); validation of *Gnawa* integration in *P. neolycopersici* (Figs. S23 and S24); *Gnawa* sequence identity and paralog divergence (Figs. S25 and S26); *Gnawa*-derived MITEs (Figs. S27 and S28); insertion preferences of gMITE and of endogenous transposable elements (Figs. S29 and S30).

**Additional file 3** (.zip). FASTA sequences. Genomic sequences of the high-confidence recent horizontal transfer events detected between plants and fungi (HGT.genes.plants.fasta) and copies of the plant-derived *Gnawa* transposon identified in the genome of *P. neolycopersici* (Gnawa.pogo.pseudoidium.fasta).

