## Additional file 2. Supplementary Figures S1-S30 with legends for "Asymmetric inter-kingdom gene transfer and a horizontal transposon invasion in a powdery mildew fungus"

### Contents


Figure S1 Maximum-likelihood phylogeny of the FAD-binding monooxygenase (FMO)

Figure S2 Maximum-likelihood phylogeny of the hypothetical protein (*HP*)

Figure S3 Maximum-likelihood phylogeny of the glycoside hydrolase (*GH*) gene

Figure S4 Maximum-likelihood phylogeny of the Oxopyrrolidines (*OP*) gene

Figure S5 Maximum-likelihood phylogeny of the glutathione S-transferase (*Fhb7*) gene

Figure S6 Maximum-likelihood phylogeny of the protein phosphatase (*PP*) gene

Figure S7 Maximum-likelihood phylogeny of a hypothetical protein (*HP*) gene

Figure S8 Maximum-likelihood phylogeny of PITH & SM-like domain (*PITH-SM*) gene

Figure S9 Maximum-likelihood phylogeny of phenolic acid decarboxylase and FAD-dependent monooxygenase (*PAD-FMO*) gene

Figure S10 Maximum-likelihood phylogeny of a hypothetical protein (*HP*) gene

Figure S11 Maximum-likelihood phylogeny of glutathione S-transferase (*GST*) gene

Figure S12 Maximum-likelihood phylogeny of the acetyltransferase (*AT*) gene

Figure S13 Maximum likelihood phylogenetic tree of *Gnawa* element displaying branch lengths

Figure S14 InterProScan annotation of transferred genes containing conserved domains

Figure S15 Chromosomal distribution of HT-derived genes across representative Pooideae genomes

Figure S16 Genomic context and flanking sequence conservation of horizontally transferred genes in *Lolium perenne*

Figure S17 RNA-seq read count of the hypothetical protein (HP) and glycoside hydrolase (GH) families in the Pooideae

Figure S18 Intraspecific copy number variation of horizontally acquired loci in *Lolium perenne*

Figure S19 Expression profiles of horizontally transferred genes in non-vascular plants

Figure S20 Structural organization and transcriptional evidence of representative HT loci in *Marchantia polymorpha*

Figure S21 Intraspecific presence/absence polymorphism of horizontal gene transfers across *Marchantia polymorpha* wild accessions

Figure S22 Phylogenetic placement of the *Gnawa* transposable element

Figure S23 Read coverage profiles of the *Gnawa* element across independent *Pseudoidium neolycopersici* genome sequencing datasets

Figure S24 PacBio long reads spanning the *Gnawa* terminal inverted repeat (TIR) junctions

Figure S25 Pairwise nucleotide sequence identity of full-length *Gnawa* homologs across different plant orders

Figure S26 Intraspecies paralog divergence of *Gnawa* homologs across host plants and *P. neolycopersici*

Figure S27 Comparative sequence alignment of *Gnawa*-derived MITEs from plants and fungi

Figure S28 Phylogenetic relationship of *Gnawa*-derived MITEs across lineages

Figure S29 Distance to nearest genes for gMITE and endogenous transposable elements (TEs) in *P. neolycopersici*

Figure S30 Insertion landscape of gMITE and of the endogenous MITE families of *P. neolycopersici* relative to random expectation

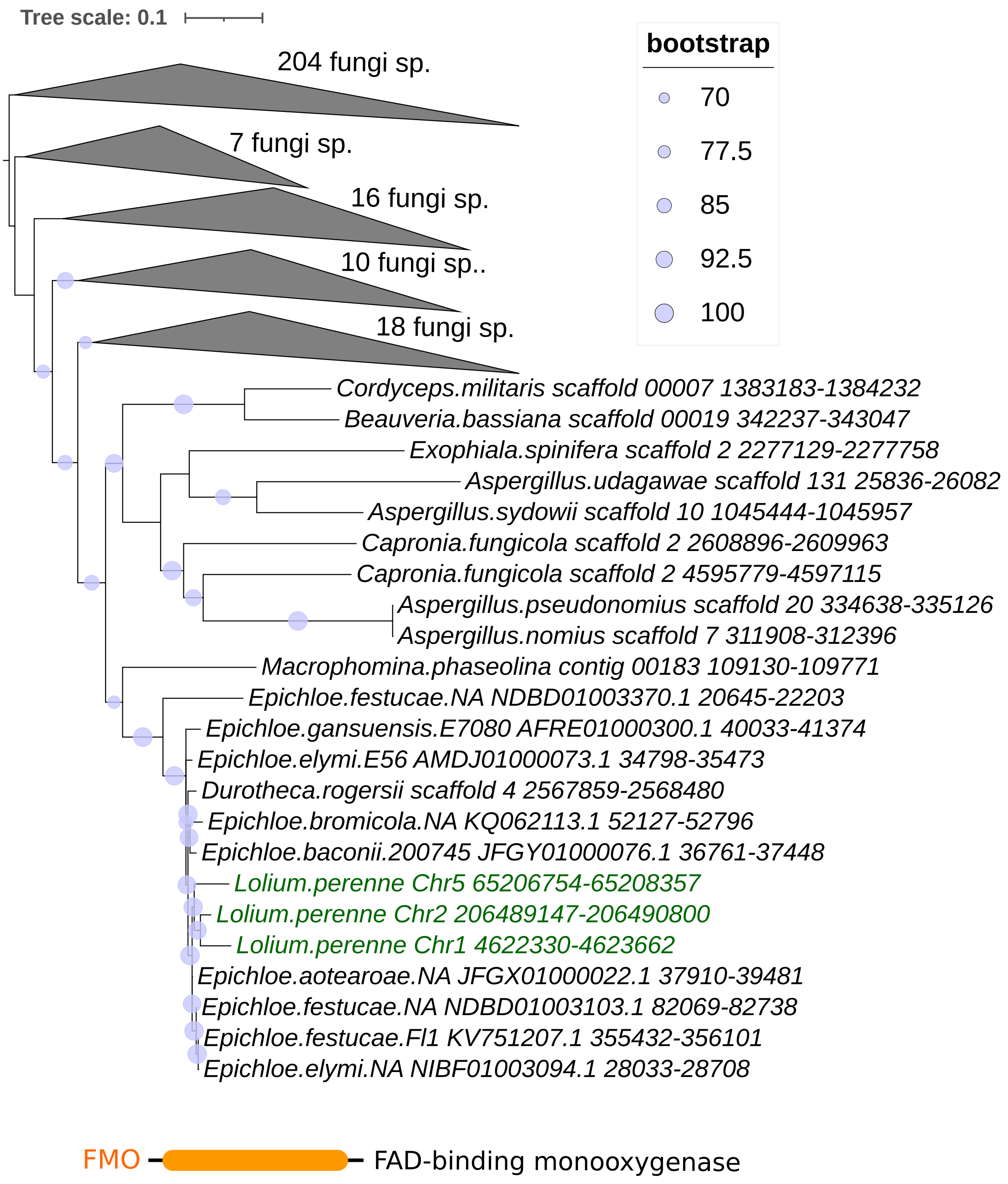


**Figure S1. Maximum-likelihood phylogeny of the FAD-binding monooxygenase (FMO).** The phylogeny was inferred from nucleotide sequences. Grey triangles represent collapsed fungal clades, with labels indicating the number of represented species. HT-derived FMO copies from *Lolium perenne* are shown in green. Circle size is proportional to ultrafast bootstrap support.

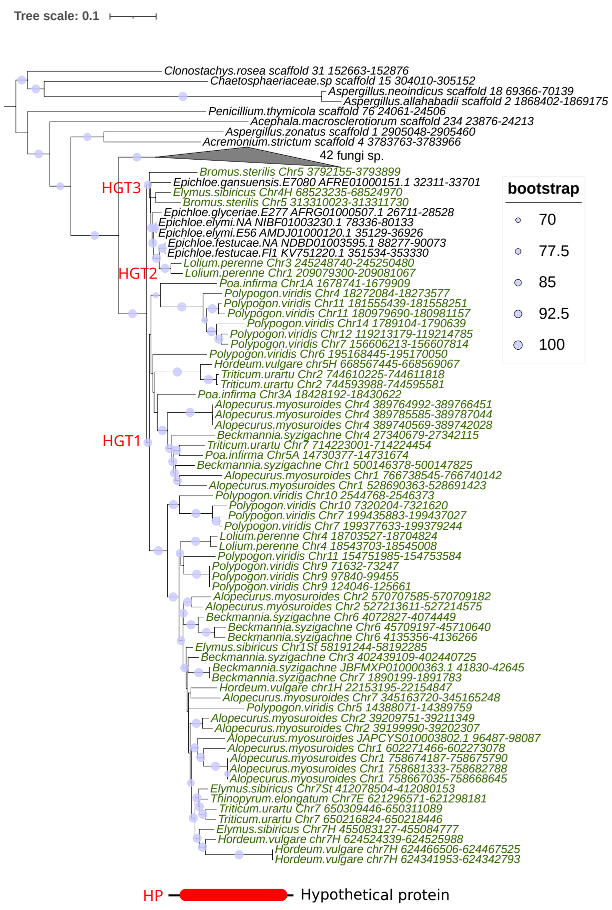


**Figure S2. Maximum-likelihood phylogeny of the hypothetical protein (*HP*).** The phylogeny was inferred from nucleotide sequences. HT-derived HP gene copies from the Pooideae species are shown in green. Circle size is proportional to ultrafast bootstrap support. HGT1 denotes the ancestral acquisition in the common ancestor of the Pooideae; HGT2 and HGT3 denote the two independent, more recent transfers.

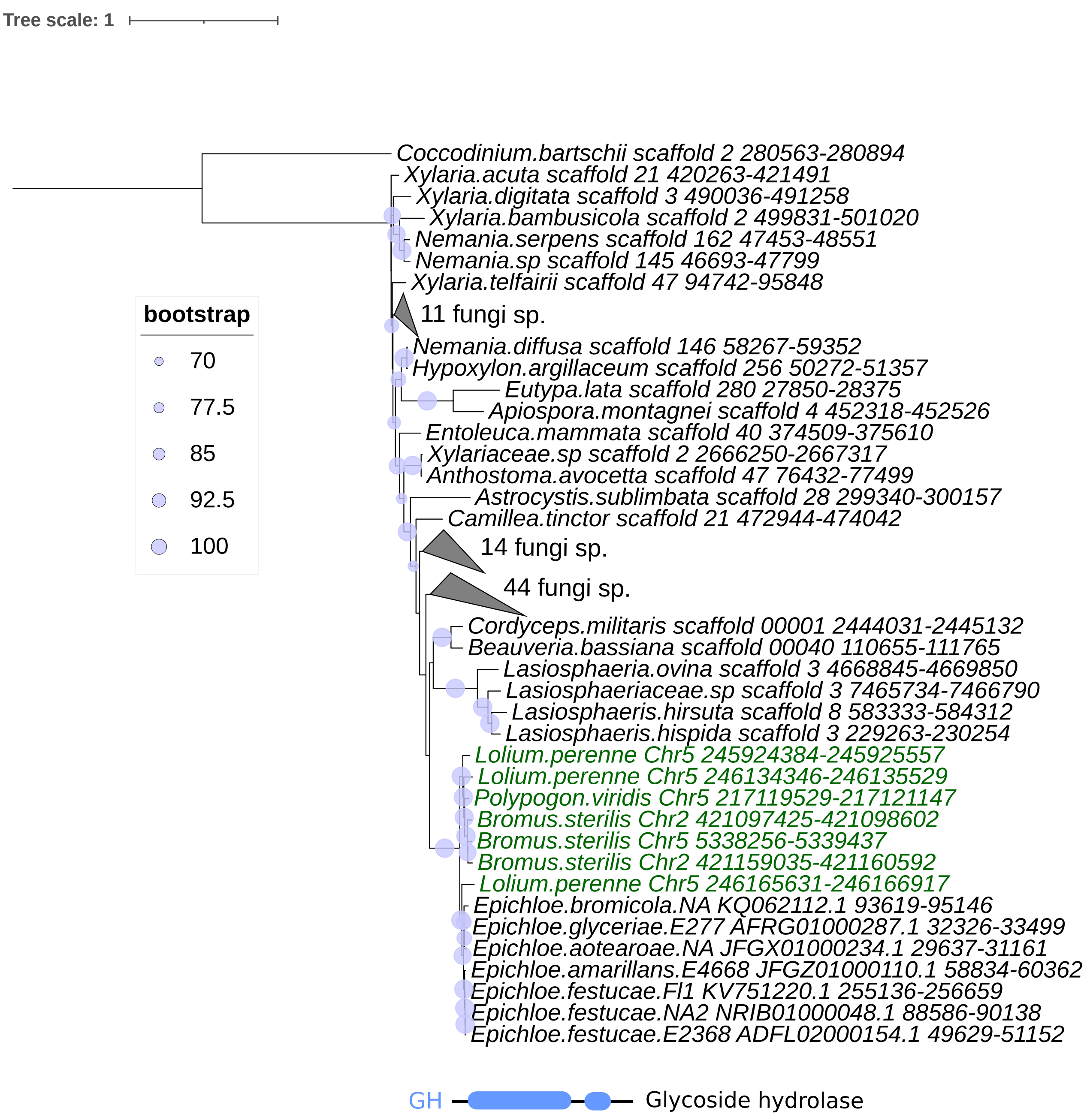


**Figure S3. Maximum-likelihood phylogeny of the glycoside hydrolase (*GH*) gene.** The phylogeny was inferred from nucleotide sequences. Grey triangles represent collapsed fungal clades, with labels indicating the number of represented species. HT-derived *GH* copies from the Pooideae species are shown in green. Circle size is proportional to ultrafast bootstrap support.

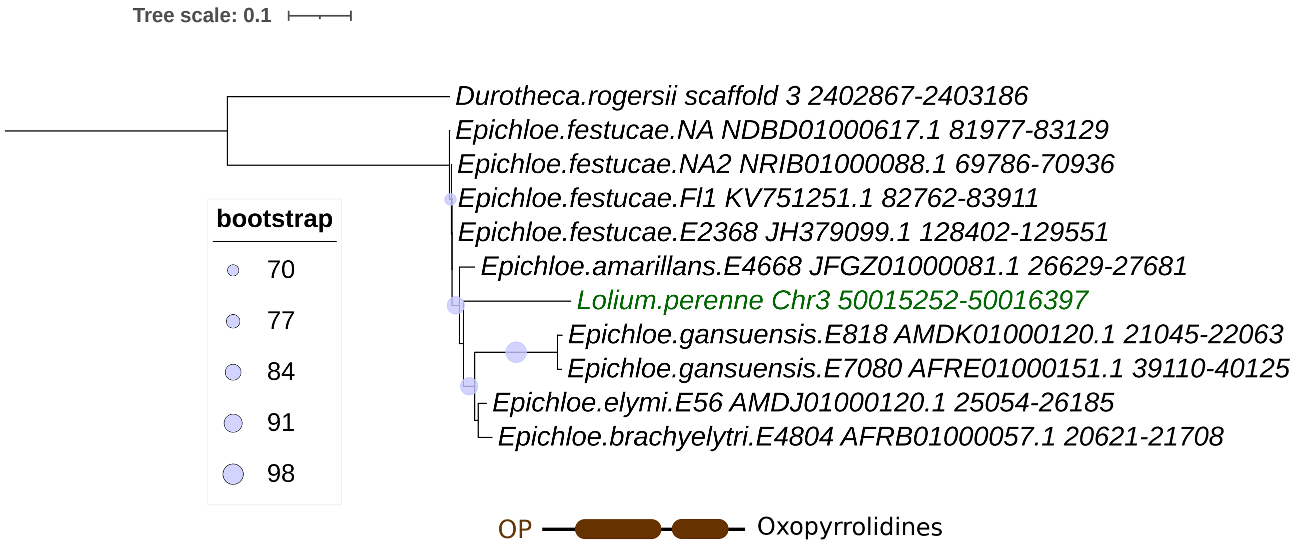


**Figure S4. Maximum-likelihood phylogeny of the Oxopyrrolidines (*OP*) gene.** The phylogeny was inferred from nucleotide sequences. *Lolium perenne* homolog is shown in green. Circle size is proportional to ultrafast bootstrap support.

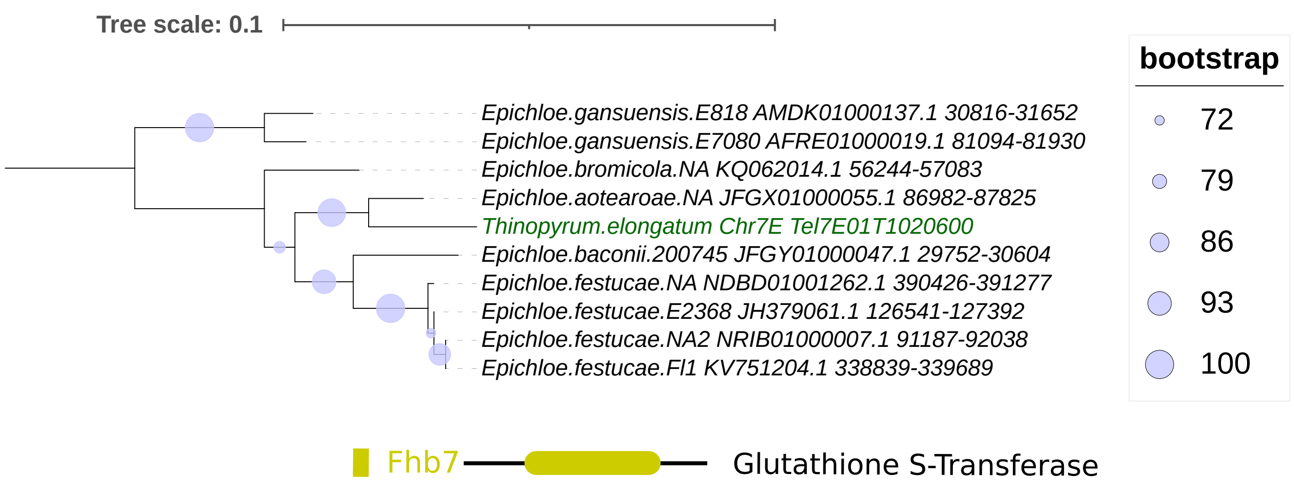


**Figure S5. Maximum-likelihood phylogeny of the glutathione S-transferase (*Fhb7*) gene.** The phylogeny was inferred from nucleotide sequences. *Thinopyrum elongatum* homolog is shown in green. Circle size is proportional to ultrafast bootstrap support.

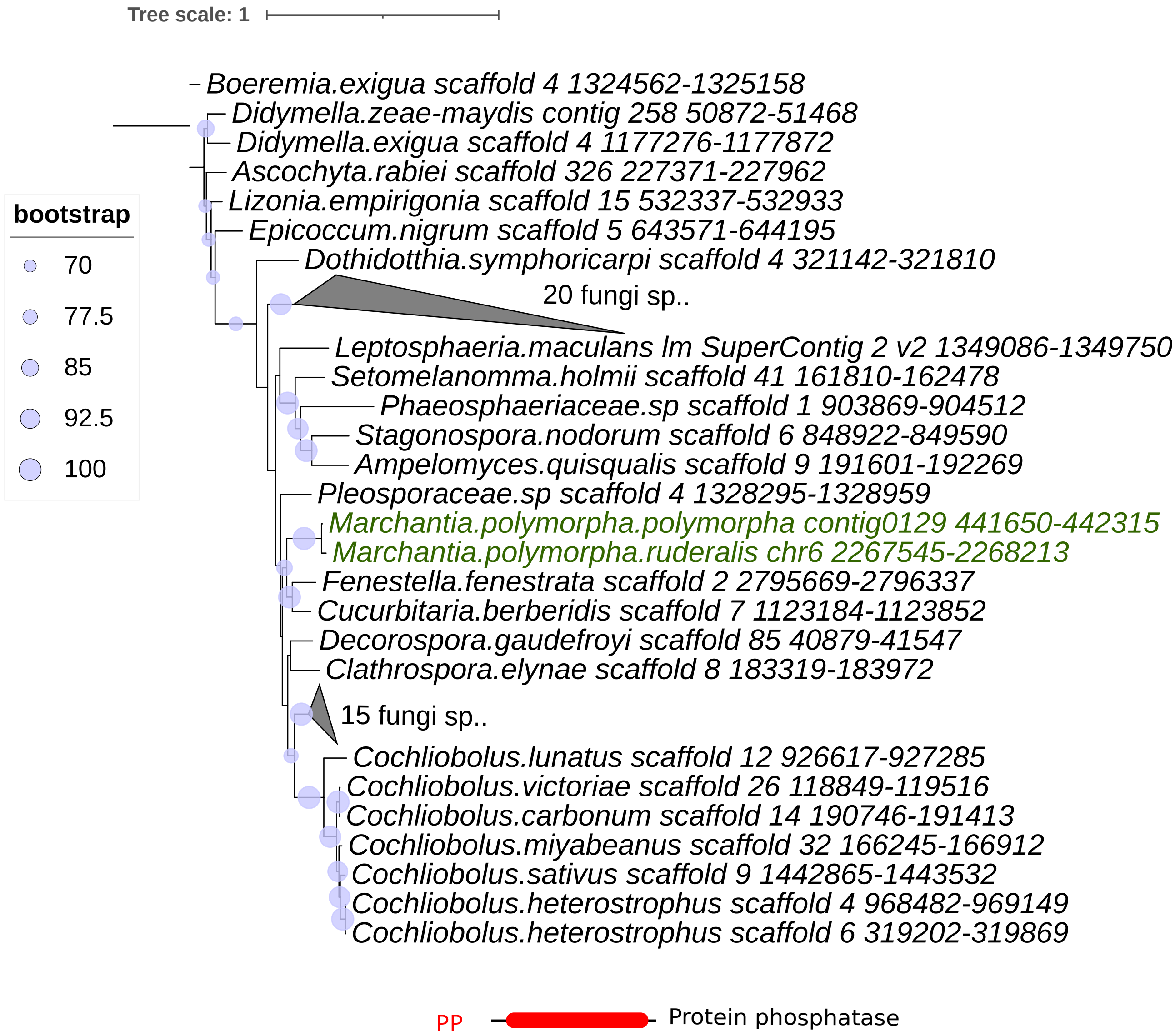


**Figure S6. Maximum-likelihood phylogeny of the protein phosphatase (*PP*) gene.** The phylogeny was inferred from nucleotide sequences. *Marchantia polymorpha* homologs are shown in green. Grey triangles represent collapsed fungal clades, with labels indicating the number of represented species. Circle size is proportional to ultrafast bootstrap support.

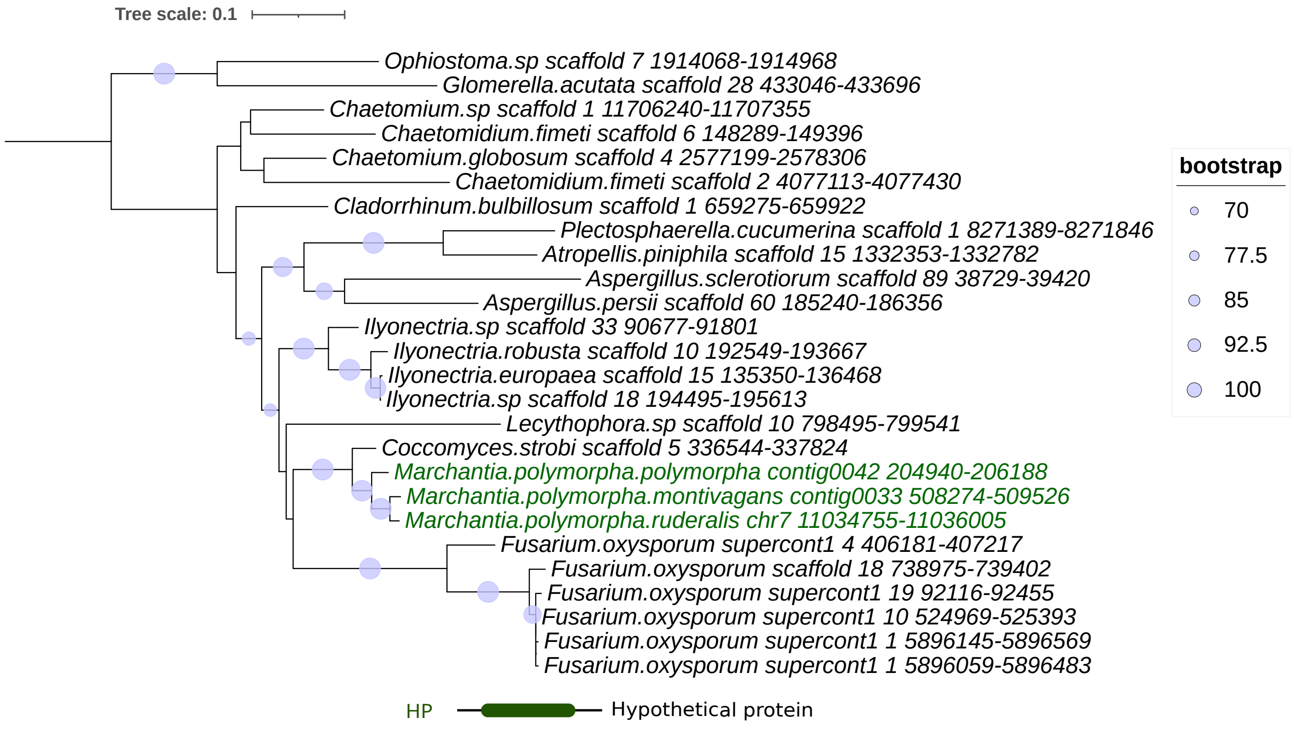


**Figure S7. Maximum-likelihood phylogeny of a hypothetical protein (*HP*) gene.** The phylogeny was inferred from nucleotide sequences. *Marchantia polymorpha* homologs are shown in green. Grey triangles represent collapsed fungal clades, with labels indicating the number of represented species. Circle size is proportional to ultrafast bootstrap support.

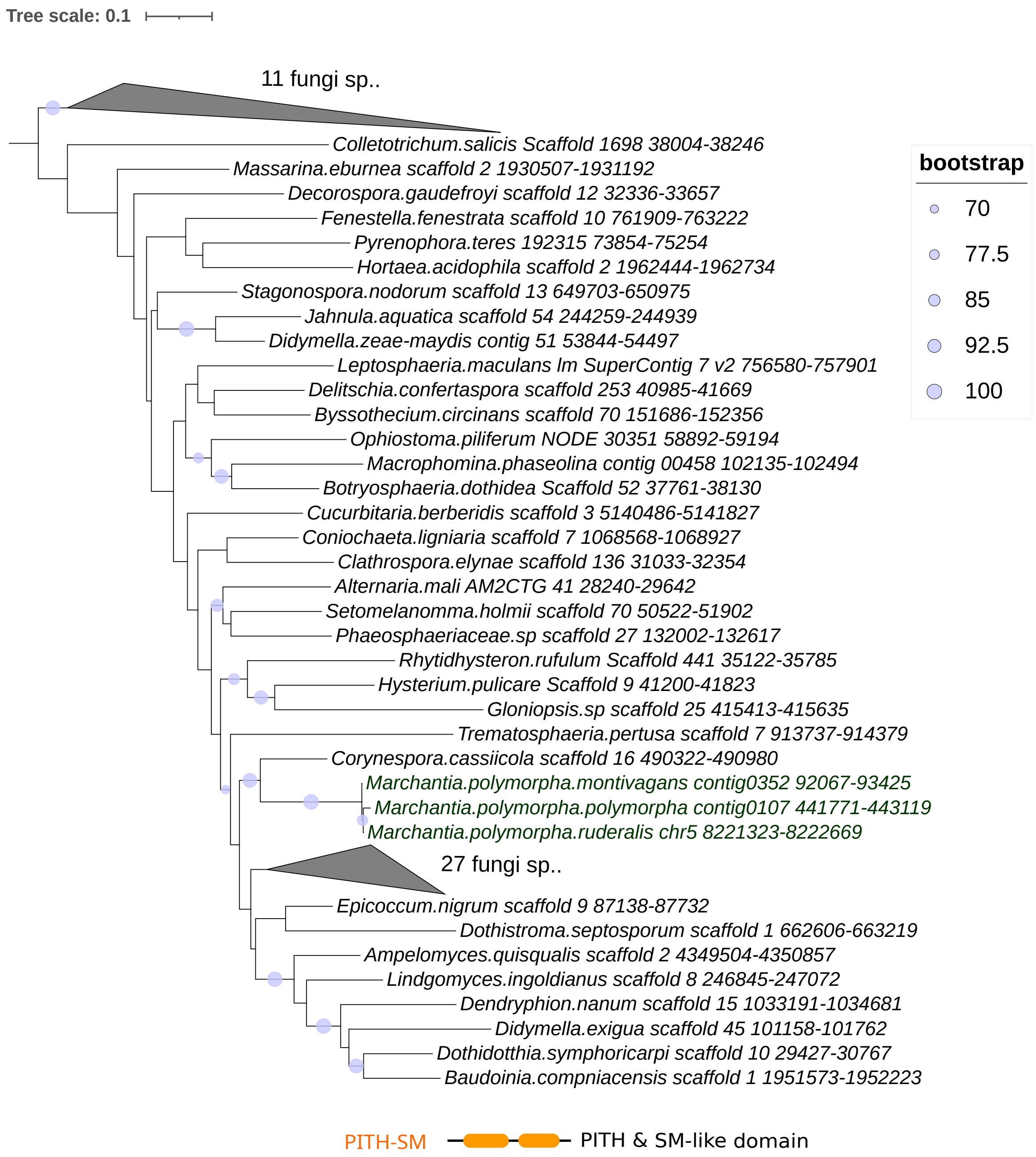


**Figure S8. Maximum-likelihood phylogeny of PITH & SM-like domain (*PITH-SM*) gene.** The phylogeny was inferred from nucleotide sequences. *Marchantia polymorpha* homologs are shown in green. Grey triangles represent collapsed fungal clades, with labels indicating the number of represented species. Circle size is proportional to ultrafast bootstrap support.

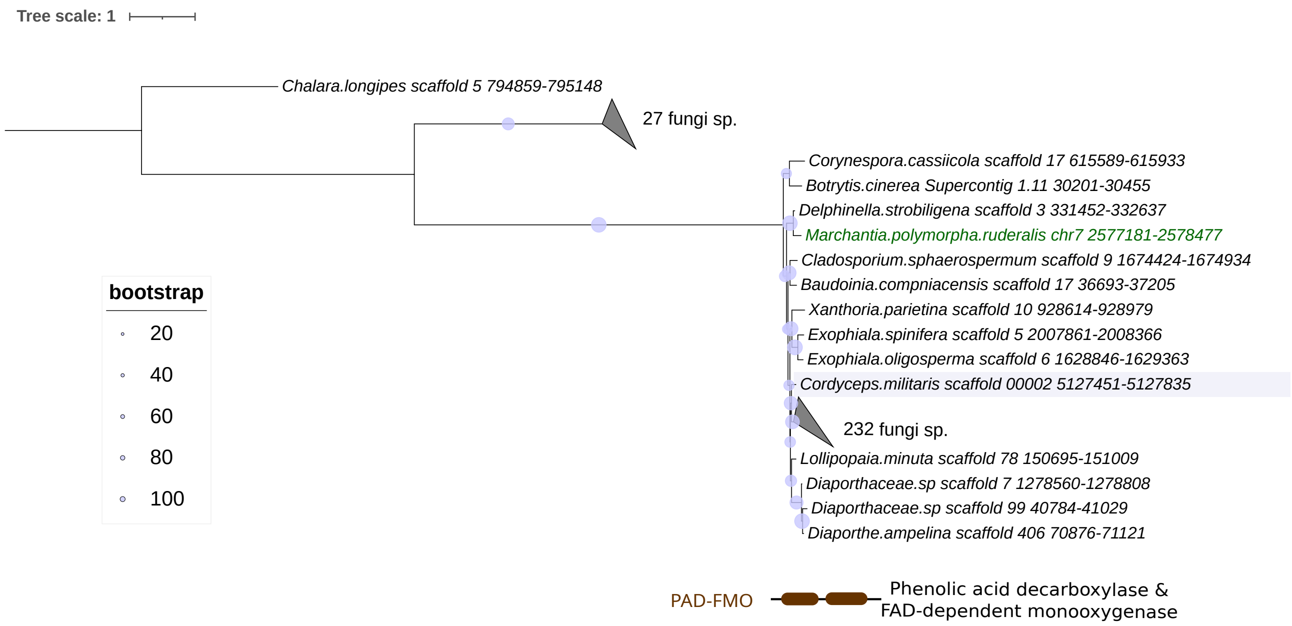


**Figure S9. Maximum-likelihood phylogeny of phenolic acid decarboxylase and FAD-dependent monooxygenase (*PAD-FMO*) gene.** The phylogeny was inferred from nucleotide sequences. *Marchantia polymorpha* homolog is shown in green. Grey triangles represent collapsed fungal clades, with labels indicating the number of represented species. Circle size is proportional to ultrafast bootstrap support.

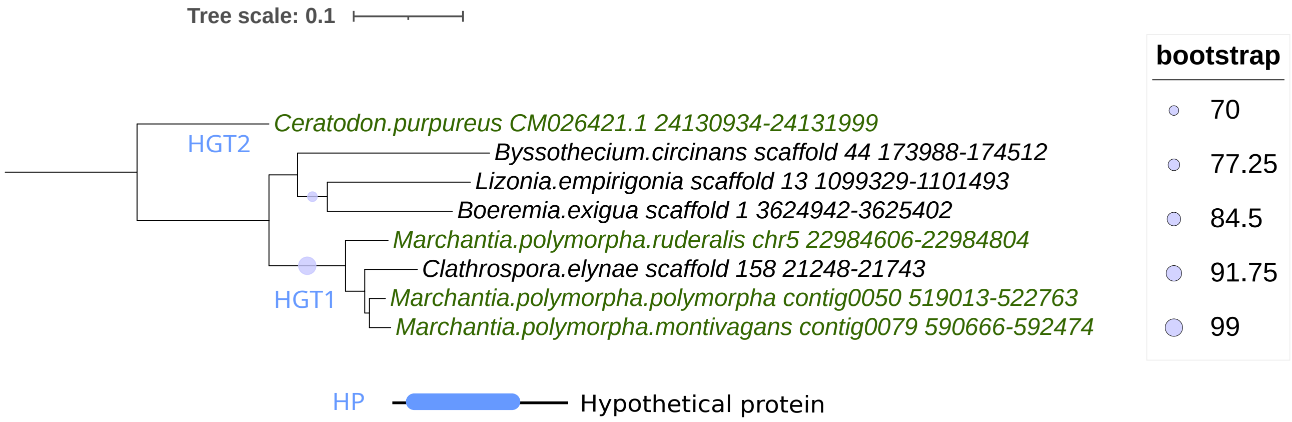


**Figure S10. Maximum-likelihood phylogeny of a hypothetical protein (*HP*) gene.** The phylogeny was inferred from nucleotide sequences. *Marchantia polymorpha* and *Ceratodon purpureus* homologs are shown in green. Circle size is proportional to ultrafast bootstrap support.

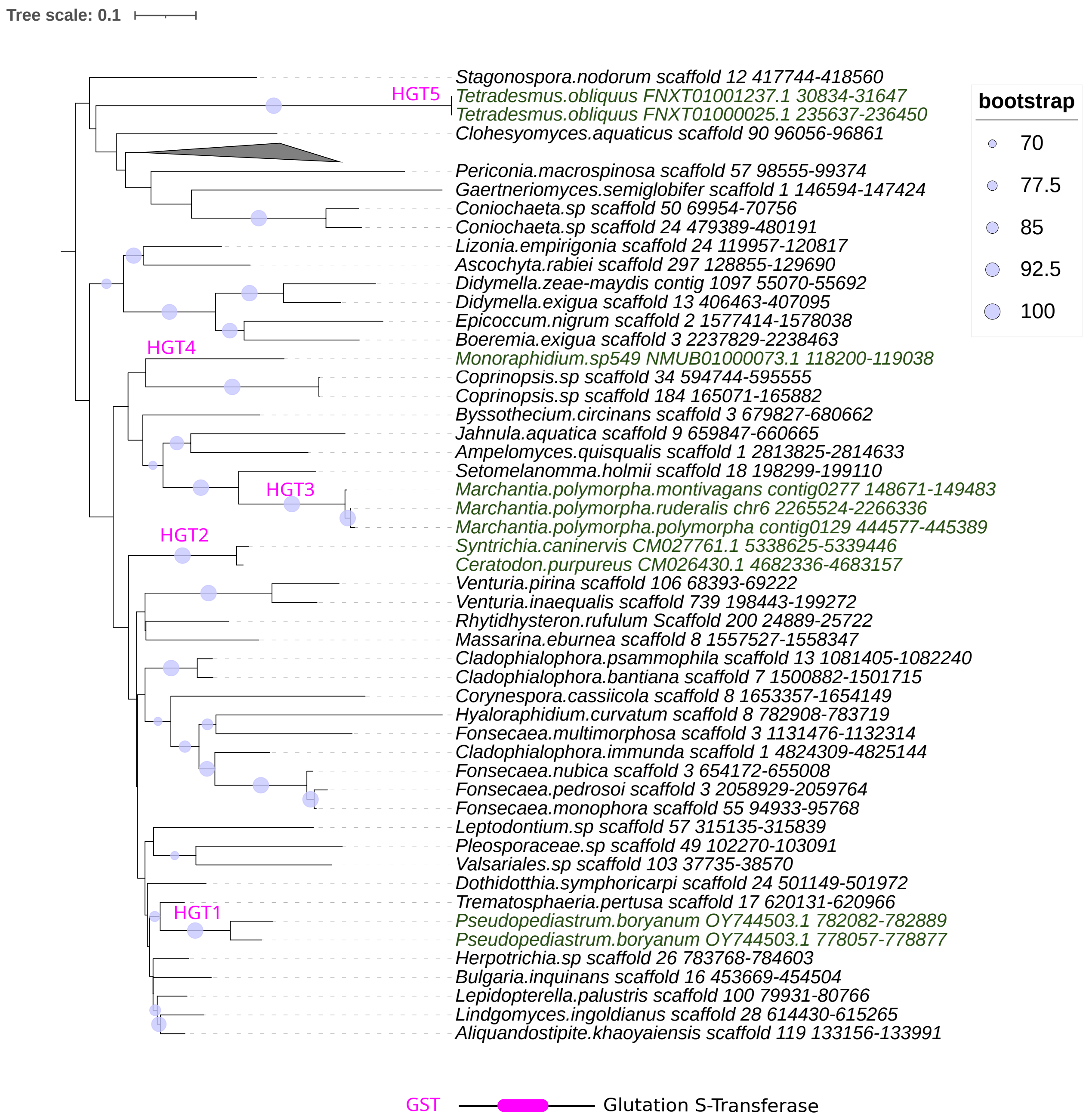


**Figure S11. Maximum-likelihood phylogeny of glutathione S-transferase (*GST*) gene.** The phylogeny was inferred from nucleotide sequences. Plant homologs are shown in green. HGT1–HGT5 indicate five independent HT events involving *GST* genes. Circle size is proportional to ultrafast bootstrap support.

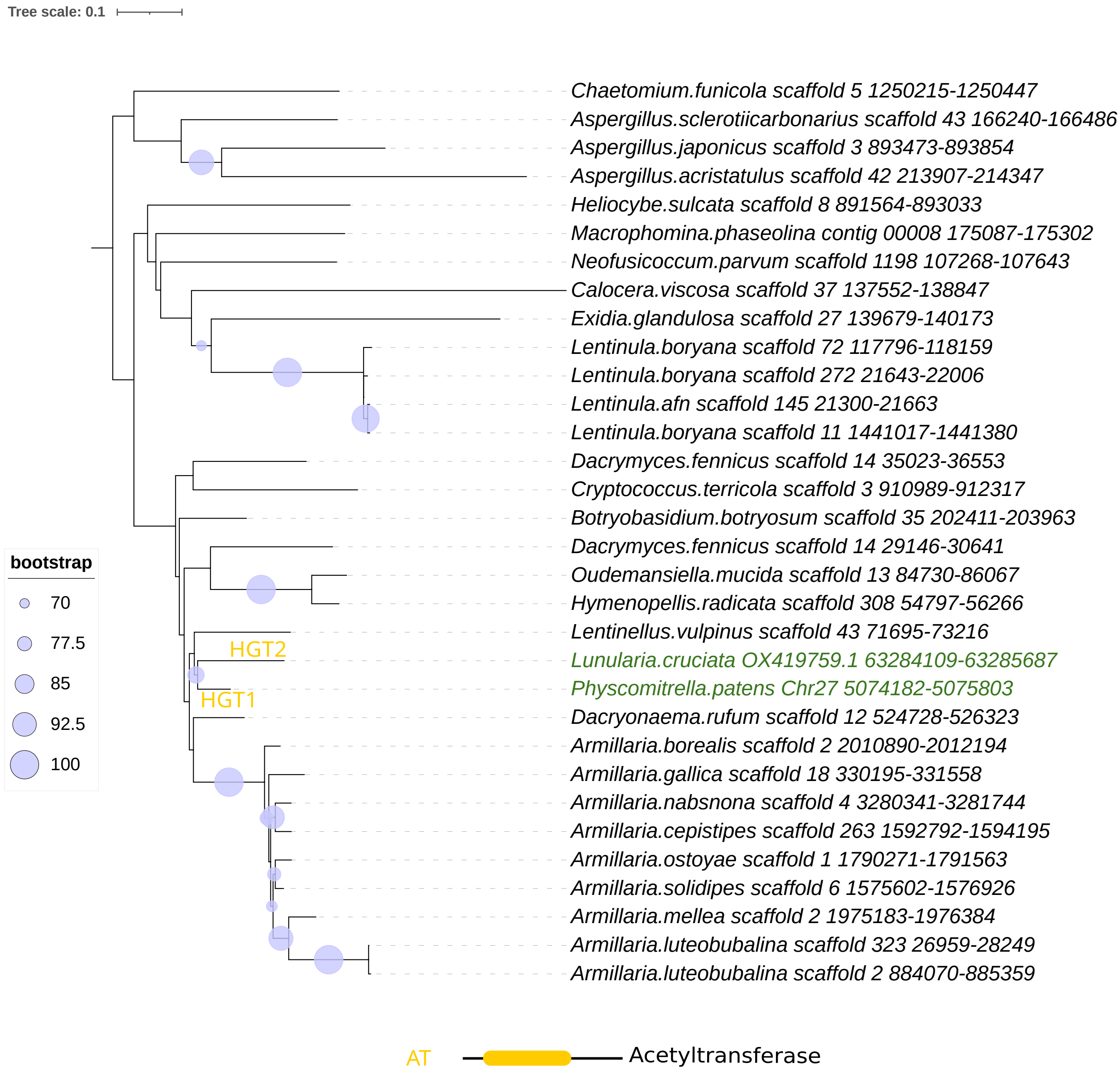


**Figure S12. Maximum-likelihood phylogeny of the acetyltransferase (*AT*) gene.** The phylogeny was inferred from nucleotide sequences. Plant homologs are shown in green. HGT1 and HGT2 indicate two independent HT events of the *AT* gene. Circle size is proportional to ultrafast bootstrap support.

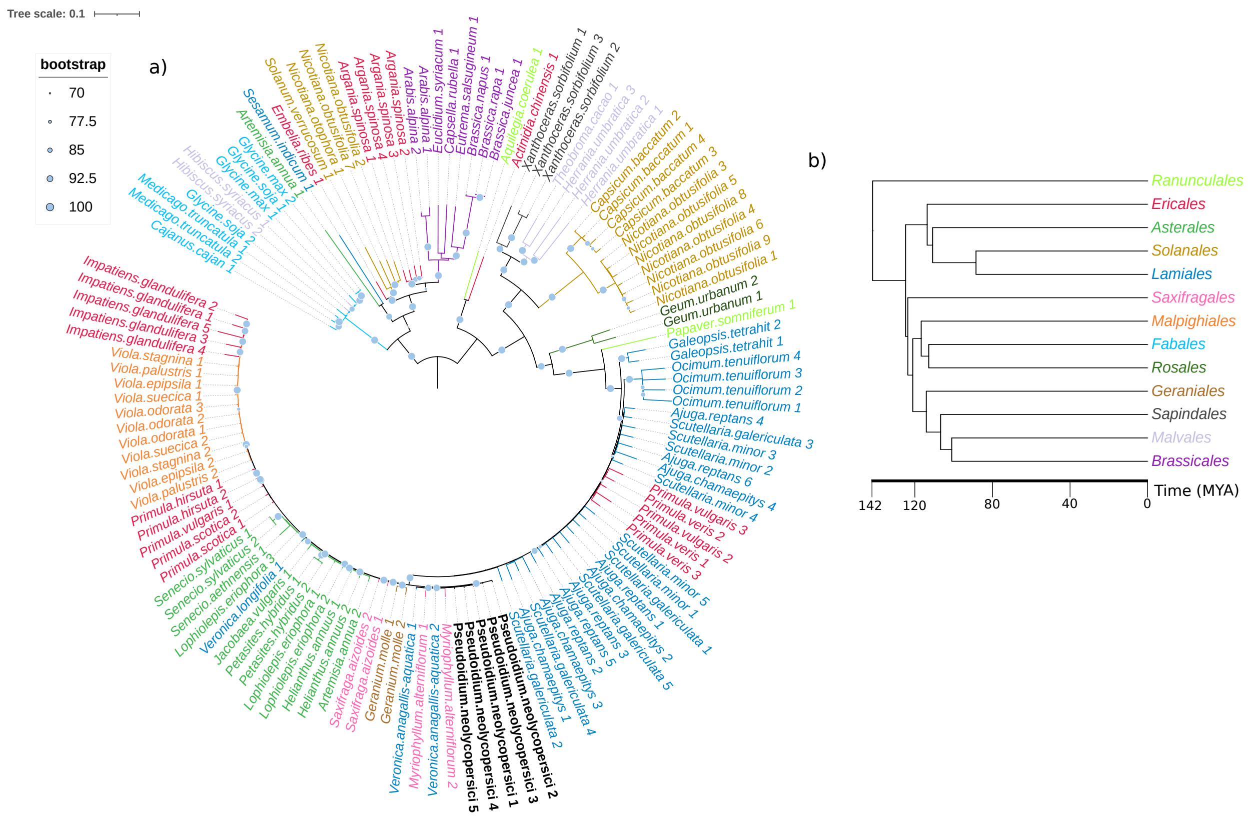


**Figure S13. Maximum likelihood phylogenetic tree of *Gnawa* element displaying branch lengths.** The phylogeny was inferred from nucleotide sequences. The unusually short branch lengths observed between *P. neolycopersici* sequences and distantly related eudicot lineages highlight the exceptionally high nucleotide identity maintained across these elements.

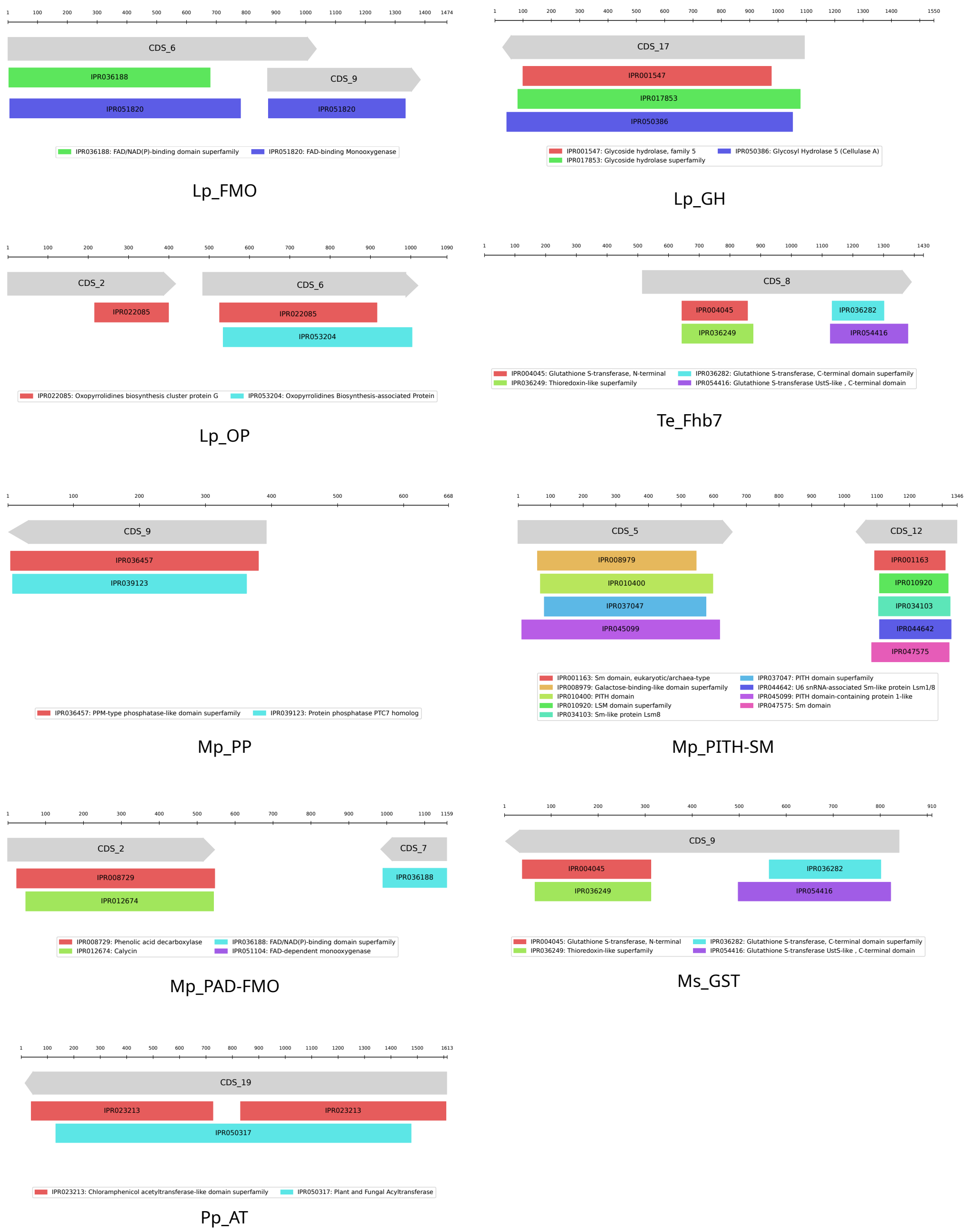


**Figure S14. InterProScan annotation of transferred genes containing conserved domains.** Transferred DNA segments were annotated for coding sequences (CDSs) and conserved domains as described in the Methods section. In each panel, the top line represents the transferred DNA segment (in nucleotides). Grey arrows indicate predicted coding sequences (CDS_nn) containing conserved domains, and colored boxes show the positions of the detected domains within each CDS. Abbreviations: Lp, *Lolium perenne*; Mp, *Marchantia polymorpha*; Te, *Thinopyrum elongatum*; Pp, *Physcomitrium patens*; Ms, *Monoraphidium* sp. 549. FMO, FAD-binding monooxygenase; HP, hypothetical protein; OP, oxopyrrolidines; GH, glycoside hydrolase; PP, protein phosphatase; PITH-SM, PITH & SM-like domain; PAD-FMO, phenolic acid decarboxylase/FAD-dependent monooxygenase; Fhb7, glutathione S-transferase Fhb7; AT, acetyltransferase.

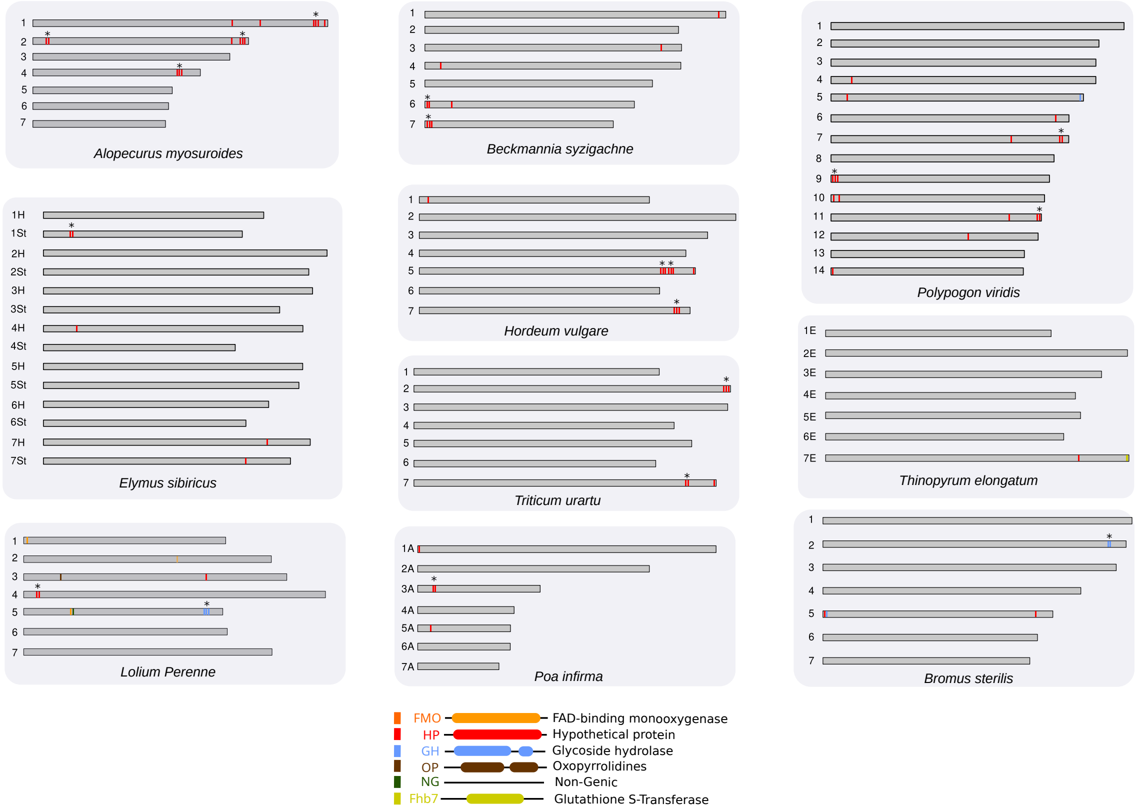


**Figure S15. Chromosomal distribution of HT-derived genes across representative Pooideae genomes.** HT-derived genes are colored according to gene family (*FMO*, FAD-binding monooxygenase; *HP*, hypothetical protein; *GH*, glycoside hydrolase; *OP*, oxopyrrolidine; *NG*, non-genic; *Fhb7*, glutathione S-transferase). Asterisks indicate tandemly duplicated copies.

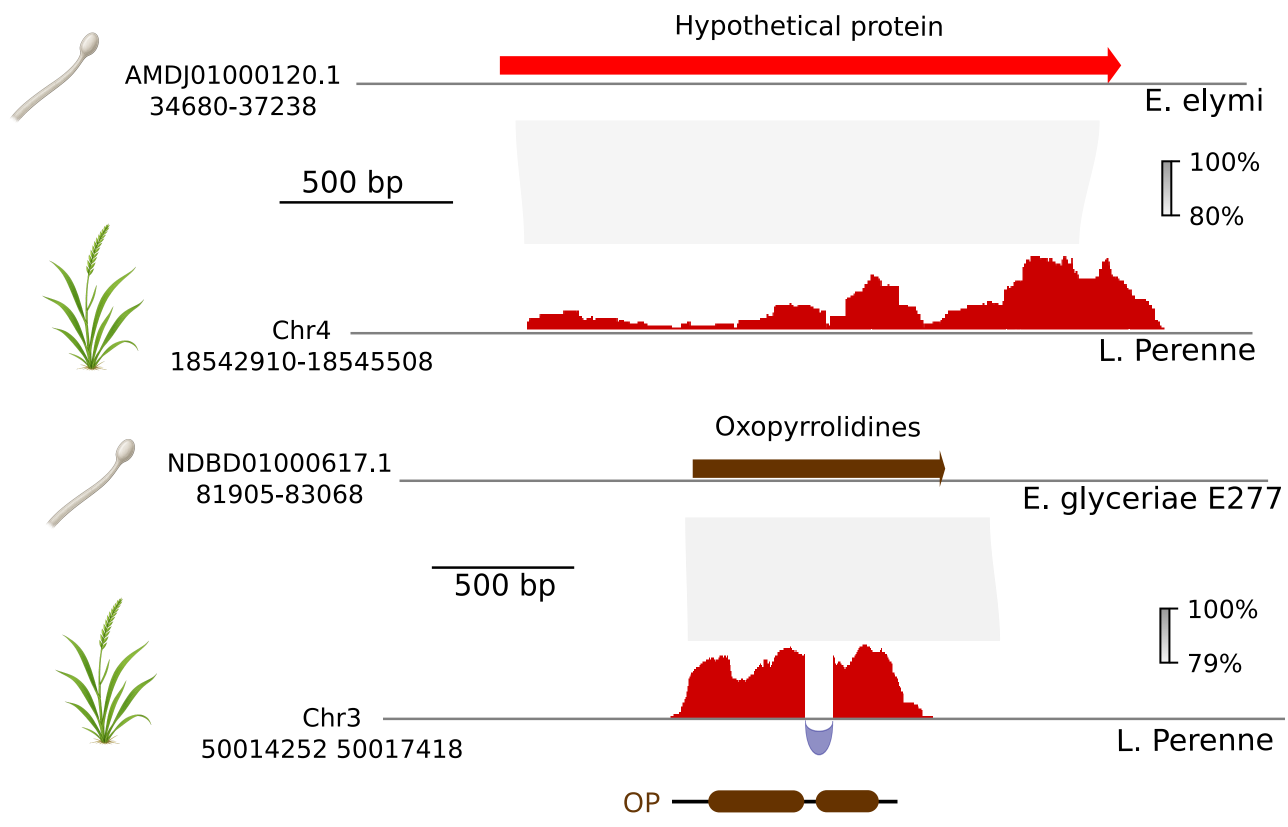


**Figure S16. Genomic context and flanking sequence conservation of horizontally transferred genes in *Lolium perenne*.** Sequence identity plots comparing *Epichloë* donor genomes to the *L. perenne* genome for the oxopyrrolidines (*OP*) and hypothetical protein (*HP*) loci. Percentage represents Blastn sequence identity.

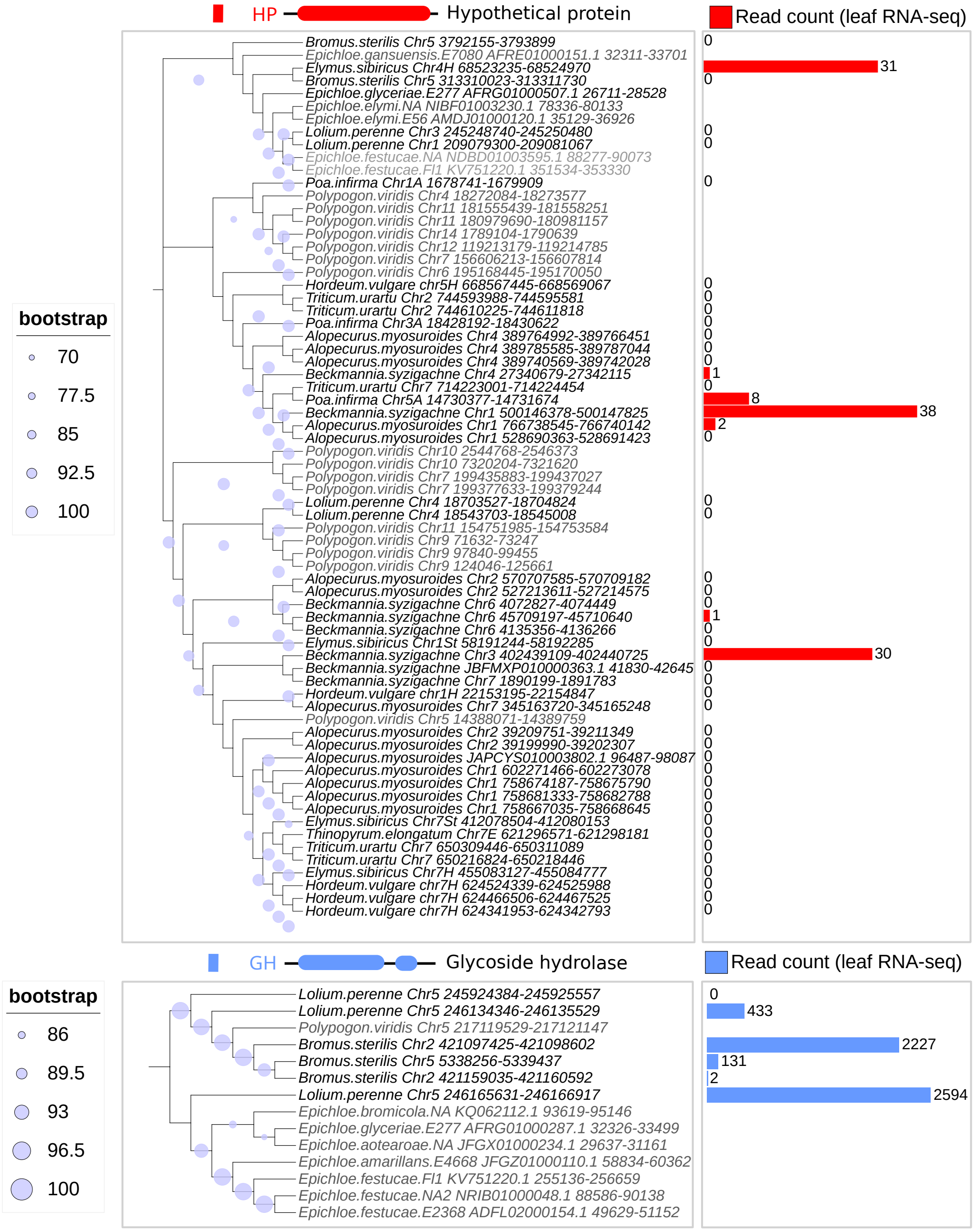


**Figure S17. RNA-seq read count of the hypothetical protein (HP) and glycoside hydrolase (GH) families in the Pooideae.** Bar plots show leaf RNA-seq read counts for the corresponding HT-derived grass homologs. Species names shown in grey indicate that no leaf RNA-seq data were available. Circle size is proportional to ultrafast bootstrap support.

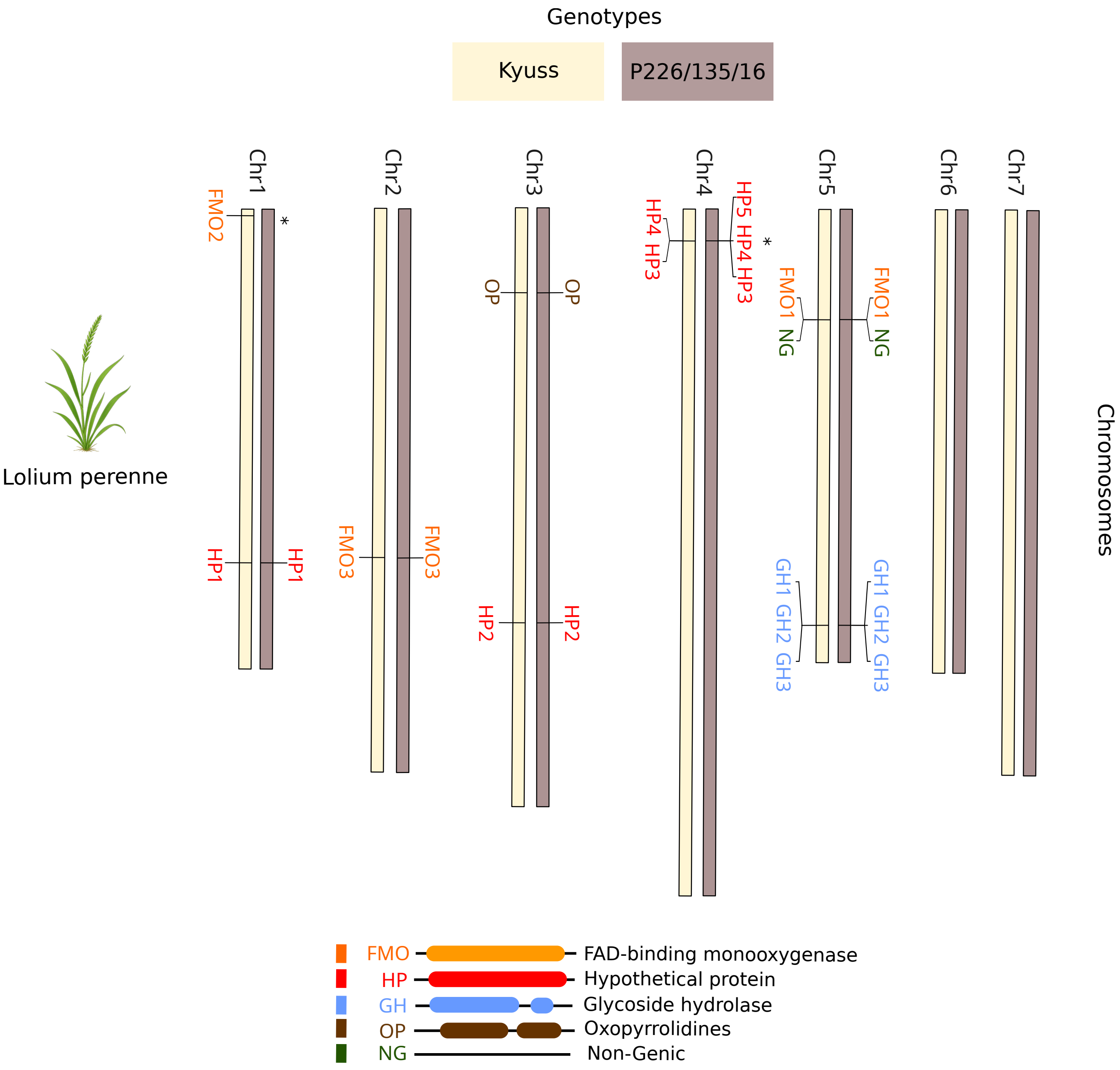


**Figure S18. Intraspecific copy number variation of horizontally acquired loci in *Lolium perenne*.** Chromosomal mapping compares the presence, absence, and tandem duplication status of the HT-derived genes (*FMO*, *GH*, *HP*, *OP*, and *NG*) between two high-quality *L. perenne* reference genomes, Kyuss and P226/135/16.

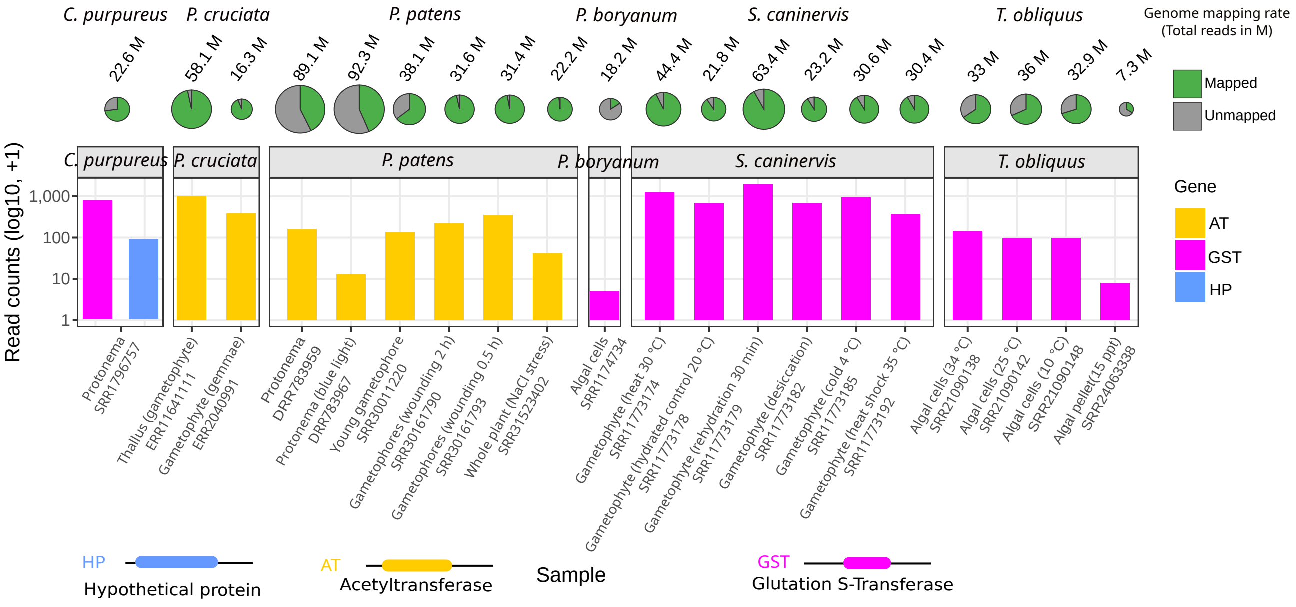


**Figure S19. Expression profiles of horizontally transferred genes in non-vascular plants.** $\mathrm{Log}_{10}$-transformed RNA-seq read counts of HT-derived genes (*HP*, hypothetical protein; *AT*, acetyltransferase; *GST*, glutathione S-transferase) across diverse tissues, developmental stages, and experimental conditions in *Ceratodon purpureus*, *Pellia cruciata*, *Physcomitrium patens*, *Syntrichia caninervis*, *Pseudopediastrum boryanum*, and *Tetradesmus obliquus*. Pie charts indicate the proportion of mapped and unmapped reads, and numbers above each chart indicate the total number of RNA-seq reads (millions).

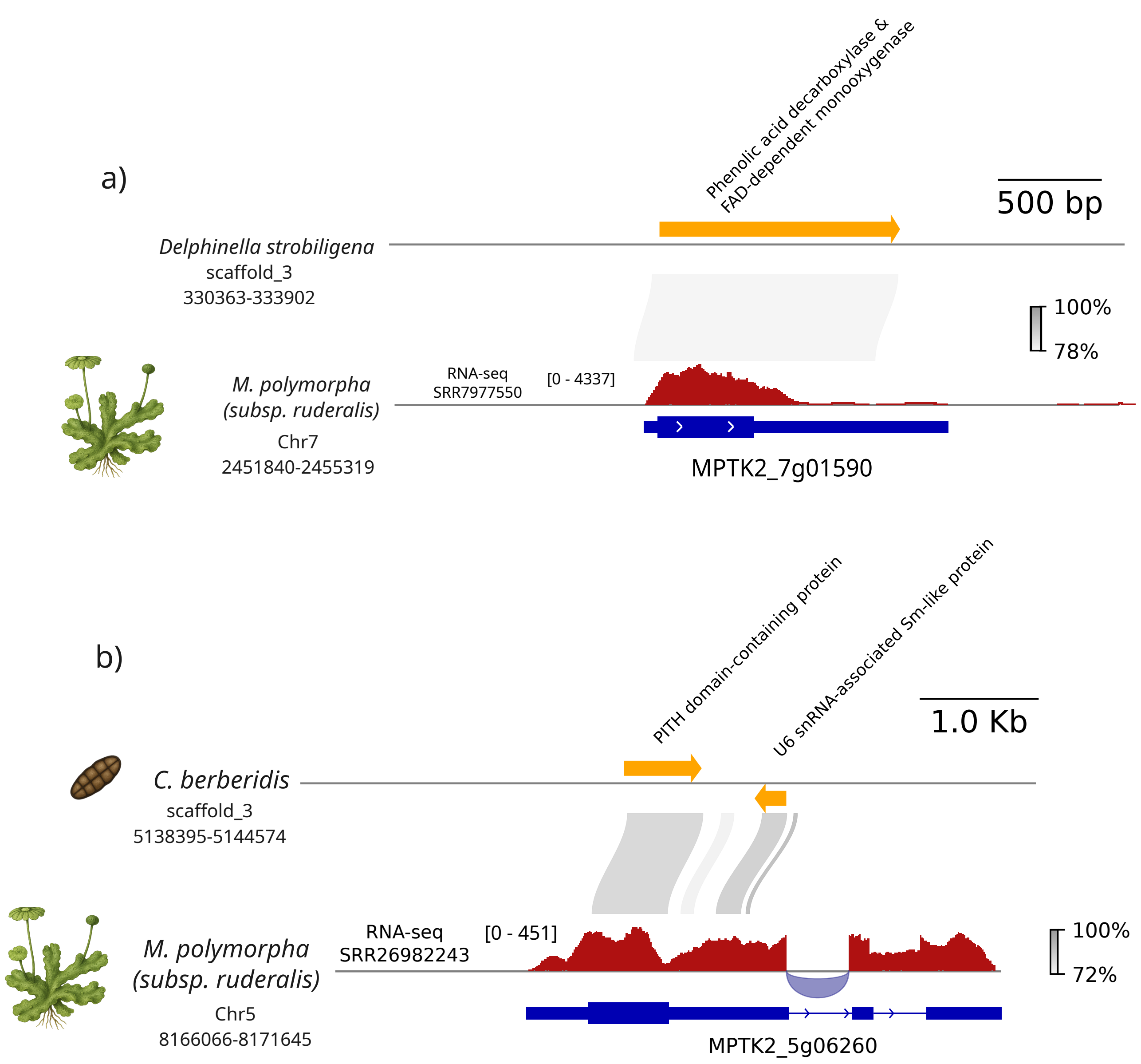


**Figure S20. Structural organization and transcriptional evidence of representative HT loci in *Marchantia polymorpha*.** **a** Genomic organization of the PAD-FMO locus showing its conservation with the putative fungal donor (*Delphinella strobiligena*) and RNA-seq read coverage across the transferred region in *M. polymorpha*. **b** Genomic organization of the *PITH-SM* locus showing its conservation with the putative fungal donor (*Corynespora cassiicola*) and its integration into a pre-existing endogenous host gene (MpTK2_5g06260), resulting in a chimeric locus.

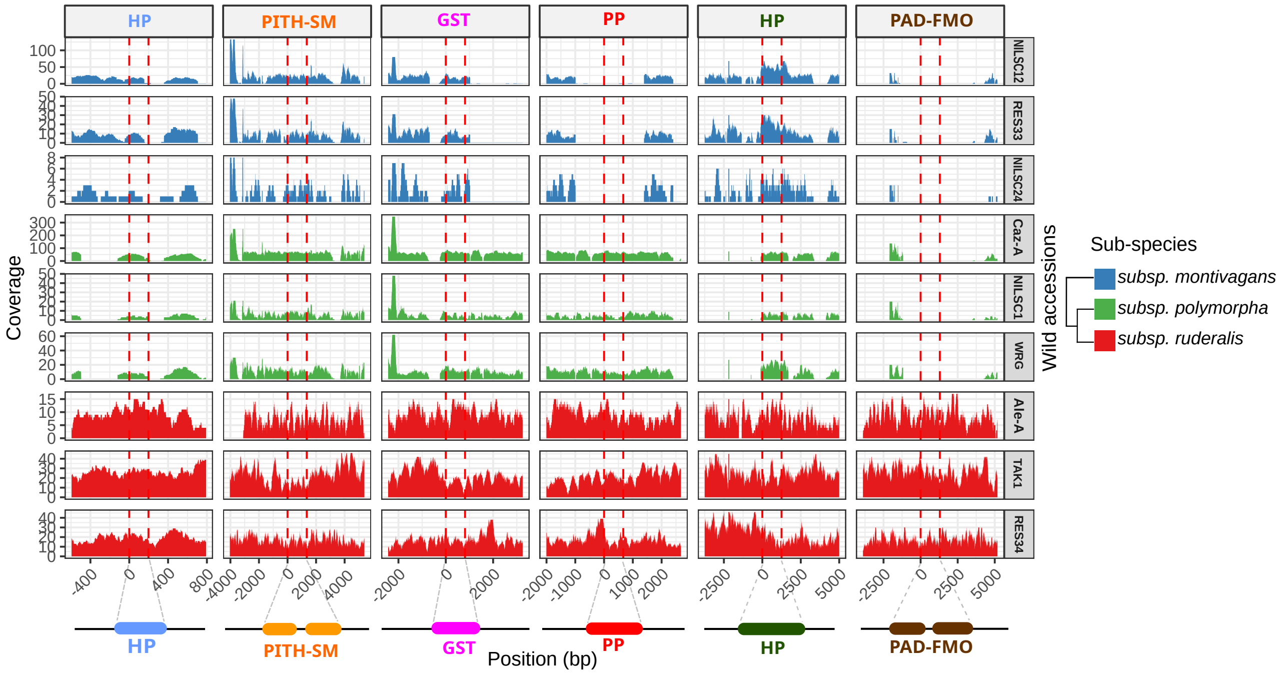


**Figure S21. Intraspecific presence/absence polymorphism of horizontal gene transfers across *Marchantia polymorpha* wild accessions.** Read coverage plots for wild accessions from the three major subspecies (*montivagans*, *polymorpha*, and *ruderalis*), confirming that differences in HT content (*HP*, *PITH-SM*, *GST*, *PP*, and *PAD-FMO*) reflect genuine biological variation and ongoing gene loss rather than assembly artifacts.

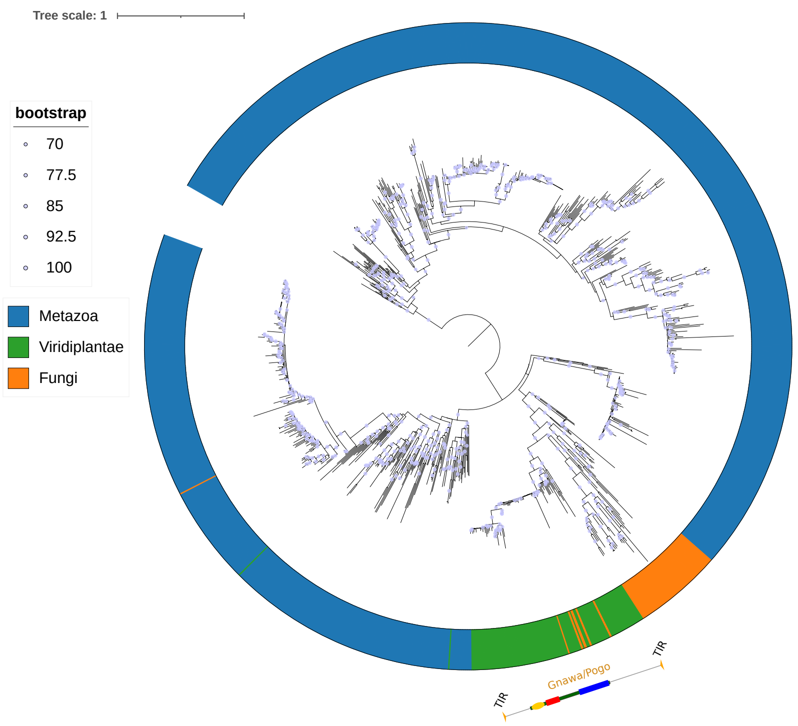


**Figure S22. Phylogenetic placement of the *Gnawa* transposable element.** Maximum likelihood tree based on the transposase domain of known *Tc1/mariner* elements, placing the *Gnawa* transposon within the *pogo*-like clade typical of Viridiplantae (plants) and distinct from typical fungal elements.

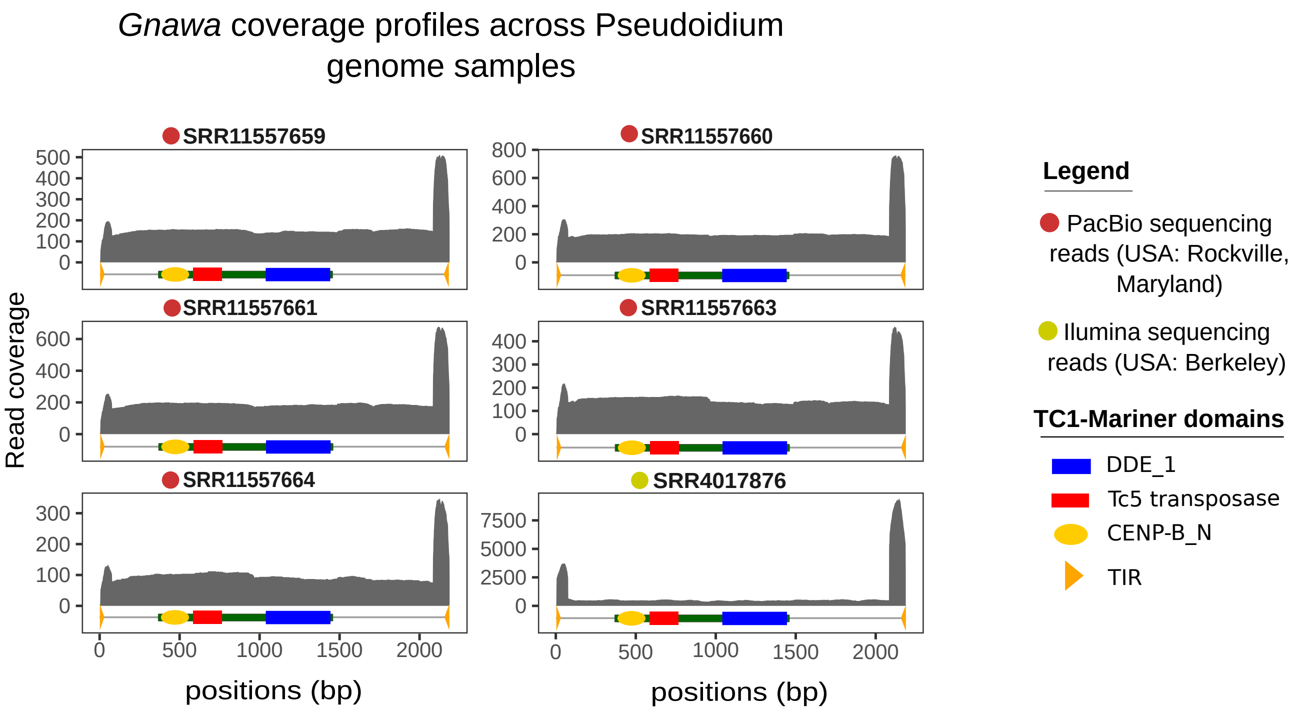


**Figure S23. Read coverage profiles of the *Gnawa* element across independent *Pseudoidium neolycopersici* genome sequencing datasets.** Whole-genome Illumina and PacBio sequencing reads from two independent *P. neolycopersici* isolates map continuously across the full-length *Gnawa* consensus sequence.

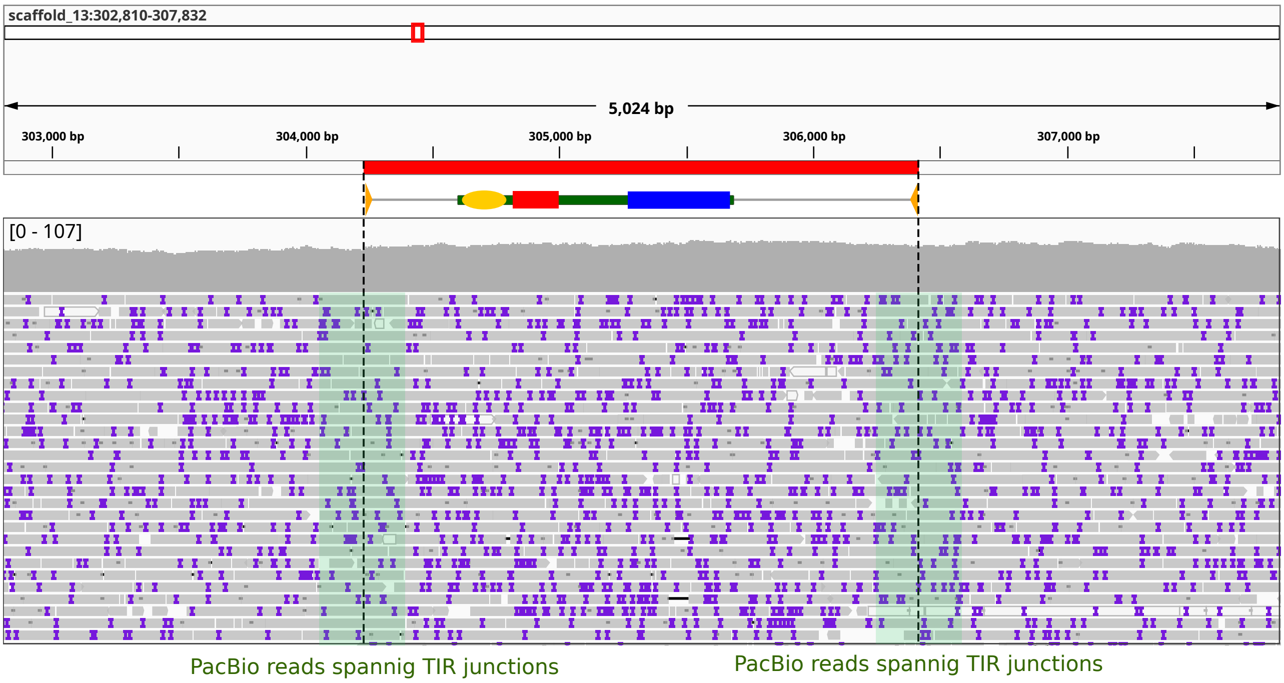


**Figure S24. PacBio long reads spanning the *Gnawa* terminal inverted repeat (TIR) junctions.** Long sequencing PacBio reads cover the entire transposon and extend into the flanking fungal genomic regions, unambiguously demonstrating the physical integration of the *Gnawa* element within the *P. neolycopersici* genome.

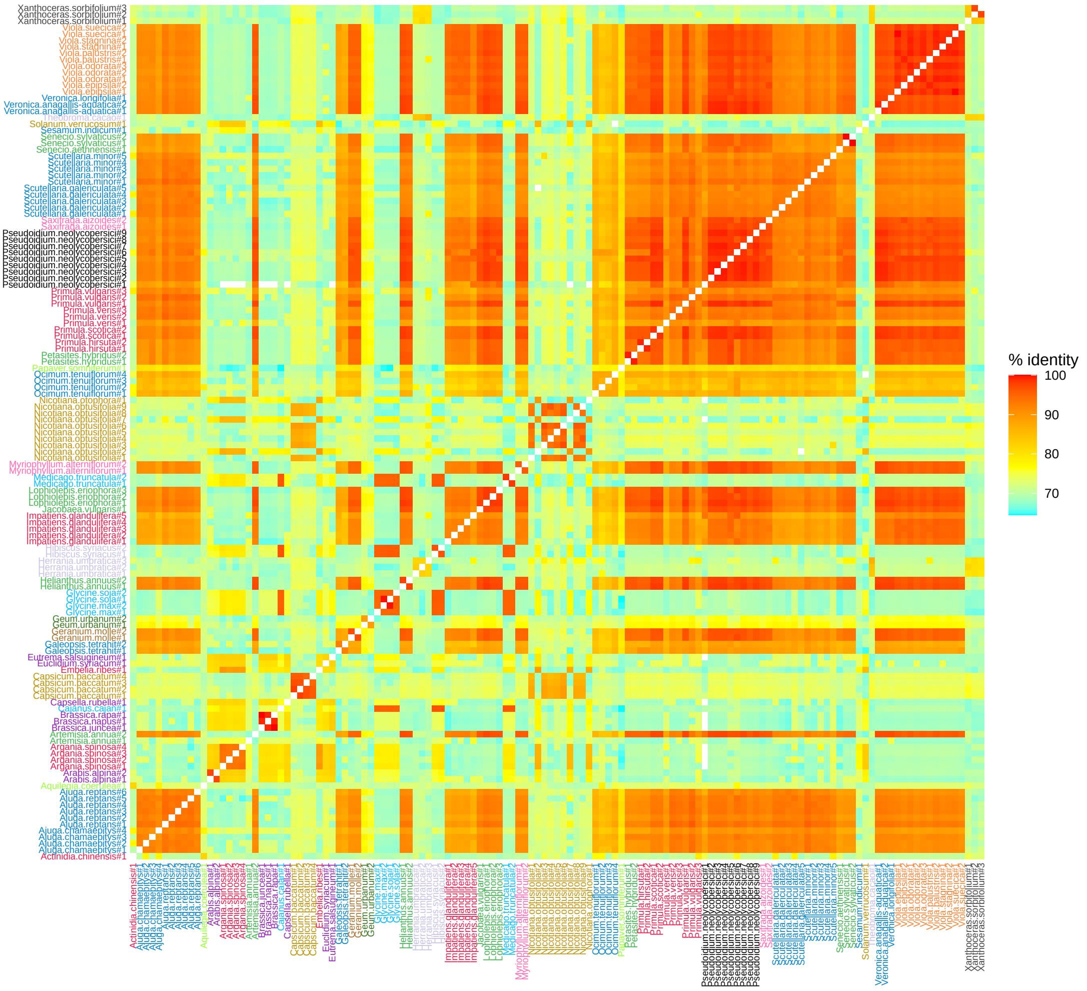


**Figure S25. Pairwise nucleotide sequence identity of full-length *Gnawa* homologs across different plant orders.** The heatmap demonstrates the remarkably high sequence conservation (frequently exceeding 95-99% identity) among *Gnawa* copies from the fungal recipient (*P. neolycopersici*) and diverse, distantly related plant hosts.

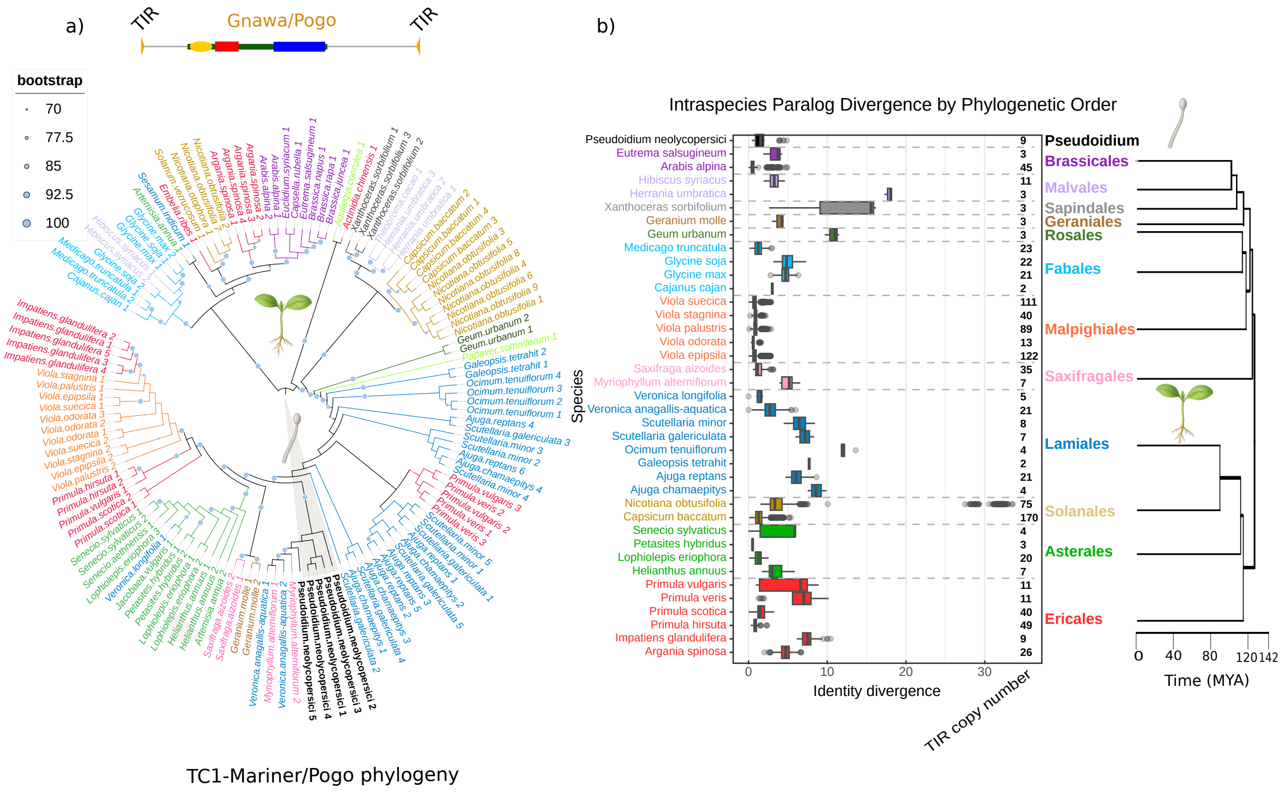


**Figure S26. Intraspecies paralog divergence of *Gnawa* homologs across host plants and *P. neolycopersici*.** Density distributions of intra-genomic sequence identity illustrate minimal divergence among *Gnawa* paralogs in several lineages, indicating recent transpositional activity and lineage-specific expansion.

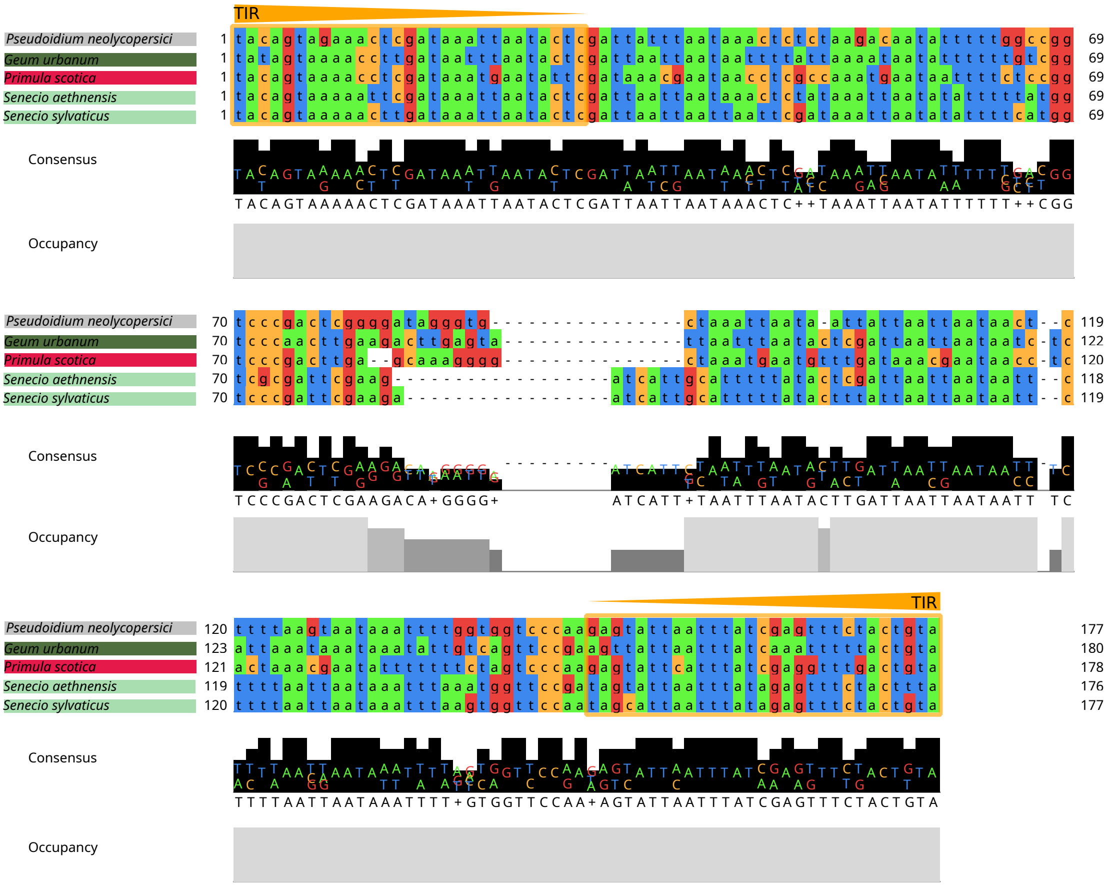


**Figure S27. Comparative sequence alignment of *Gnawa*-derived MITEs from plants and fungi.** The alignment compares the internal structures of MITEs identified in *Pseudoidium neolycopersici* and in the plant species *Geum urbanum*, *Primula scotica*, and *Senecio* spp. Terminal inverted repeats (TIRs) are highly conserved, whereas the internal regions differ substantially among lineages, consistent with independent MITE formation events.

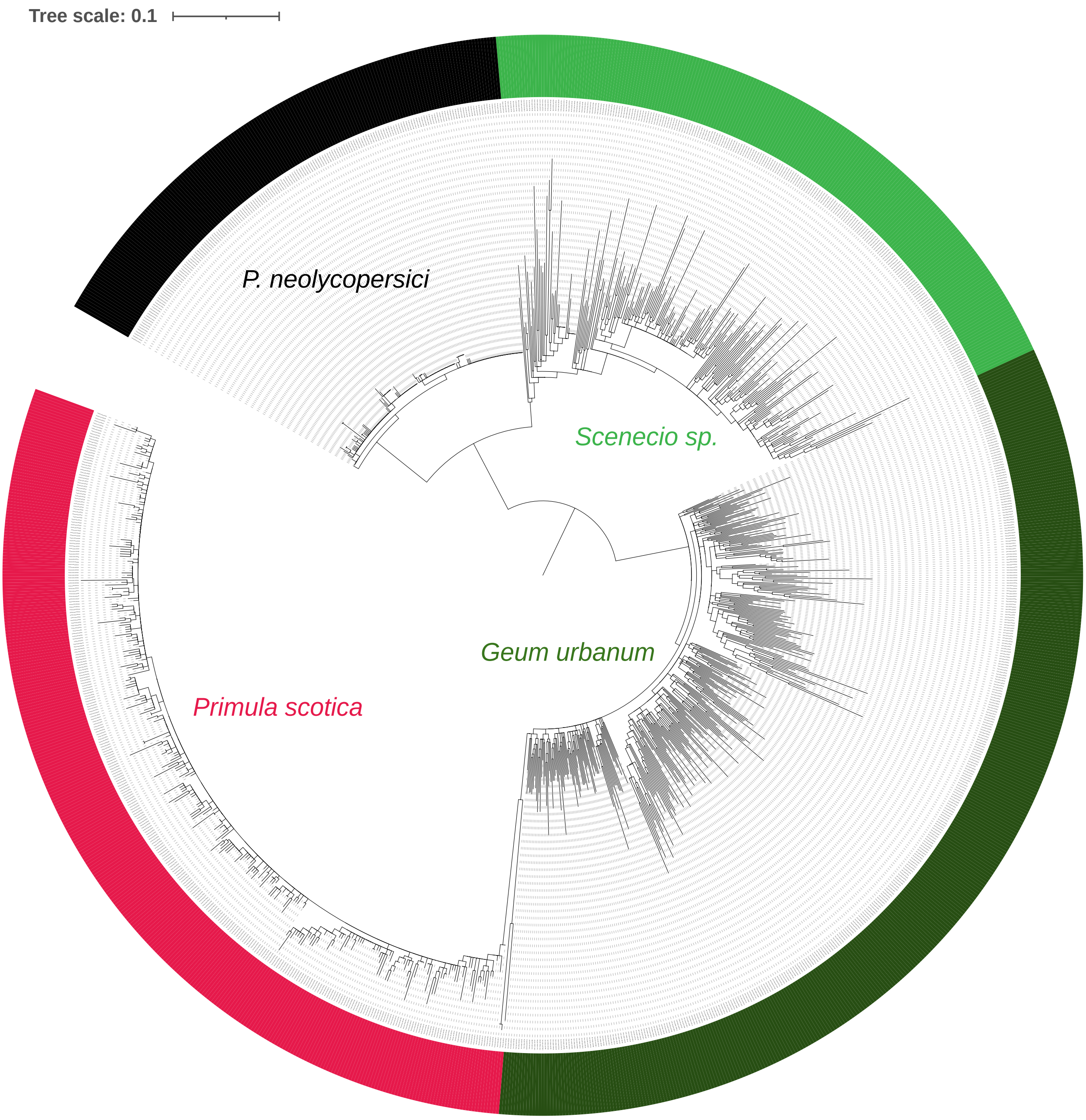


**Figure S28. Phylogenetic relationship of *Gnawa*-derived MITEs across lineages.** The phylogenetic tree includes all MITE paralogs from plants and *P. neolycopersici*.

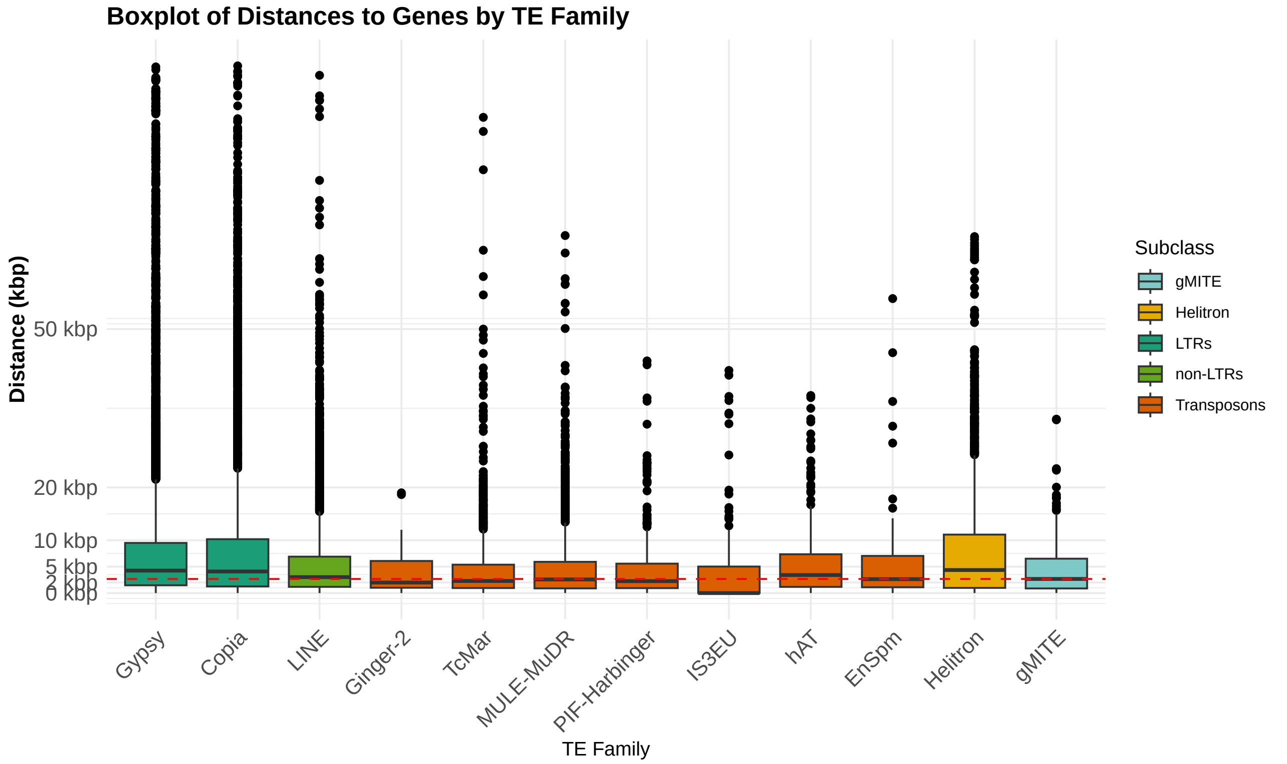


**Figure S29. Distance to nearest genes for gMITE and endogenous transposable elements (TEs) in *P. neolycopersici*.** The boxplot compares the insertion proximity to genes for gMITE against the broader TE landscape, categorized by subclass into Class I retrotransposons (LTRs, non-LTRs) and other Class II DNA transposons. Elements length differs by an order of magnitude among families and is controlled for in the permutation analysis (Additional file 1: Table S8).

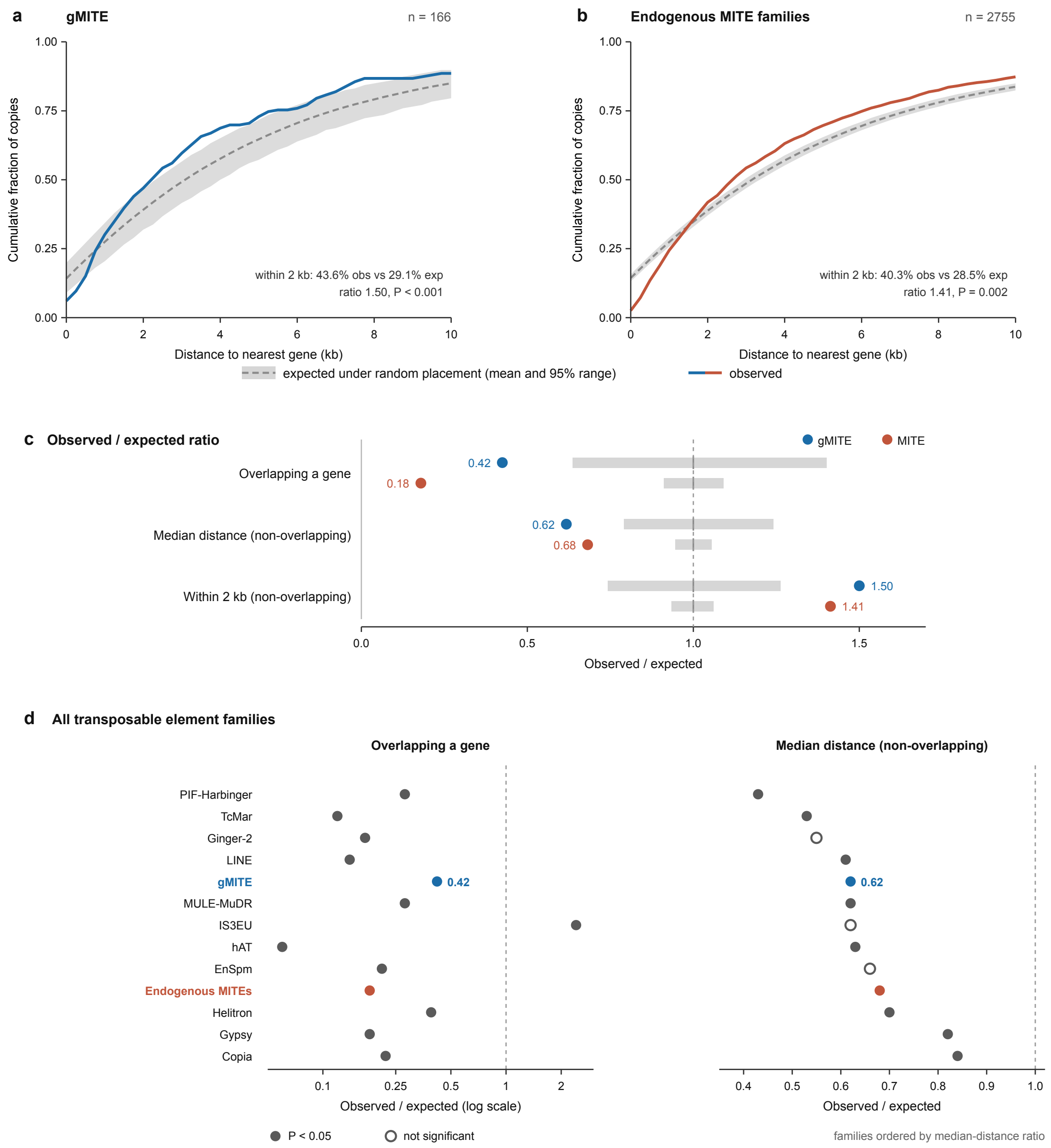


**Figure S30. Insertion landscape of gMITE and of the endogenous MITE families of *P. neolycopersici* relative to random expectation.** a, b, Cumulative distribution of the distance separating each element from the nearest annotated gene, for the 166 gMITE copies (a) and the 2,755 copies belonging to the endogenous MITE families (b). Coloured lines show the observed distribution; the grey band and dashed line show the distribution expected under random placement, obtained by repositioning every element uniformly at random within its own scaffold while preserving its length (mean and 95% range across 3,000 permutations for gMITE and 1,200 for the endogenous families). c, Ratio of observed to expected values for three statistics: the proportion of elements directly overlapping a gene, and, among elements that do not overlap a gene, the median distance to the nearest gene and the proportion lying within 2 kb of a gene. Grey bars give the 95% range of the permutation distribution, centred on a ratio of 1 (dashed line). d, The same two ratios for all thirteen transposable element categories, ordered by median-distance ratio. Filled circles denote P < 0.05; open circles denote non-significant deviations, all in families with fewer than 60 copies. Every family is depleted for insertions overlapping a gene and lies closer to genes than expected by chance, with gMITE ranking fifth of thirteen.
